# Neural and sensory basis of homing behavior in the invasive cane toad, *Rhinella marina*

**DOI:** 10.1101/2024.06.25.600658

**Authors:** Daniel A. Shaykevich, Daniela Pareja-Mejía, Chloe Golde, Andrius Pašukonis, Lauren A. O’Connell

## Abstract

The behavioral, sensory, and neural bases of vertebrate navigation are primarily described in mammals and birds. While many studies have explored amphibian navigation, none have characterized brain activity associated with navigation in the wild. To address this knowledge gap, we conducted a study on navigation in the cane toad, *Rhinella marina*. First, we performed a translocation experiment to describe how invasive cane toads in Hawaiʻi navigate home and observed homing following displacements of up to one kilometer. Next, we tested the effect of olfactory and magnetosensory manipulations on homing, as these senses are most commonly associated with amphibian navigation. We found that neither ablation alone prevents homing, further supporting that toad navigation is multimodal. Finally, we tested the hypothesis that the medial pallium, the amphibian homolog to the hippocampus, is involved in homing. Our comparisons of neural activity revealed evidence supporting a conservation of neural structures associated with navigation across vertebrates consistent with neural models of amphibian spatial cognition from recent laboratory studies. Our work furthers our evolutionary understanding of spatial behavior and cognition in vertebrates and lays a foundation for studying the behavioral, sensory, and neural bases of navigation in an invasive amphibian.

## Introduction

The ability to process spatial information is fundamental to survival for many animals, who must navigate to locate shelter, forage, and mate. Much of our understanding of spatial navigation comes from research in mammals, where the hippocampus is a key neural center for spatial representations [1,2]. In mammals [3,4] and birds [5,6], the spatial function of the hippocampus described in laboratory studies has been translated to its role in navigation in nature. In fish [7,8], reptiles [9], and amphibians [10–13], laboratory studies suggest that neural structures homologous to the hippocampus support spatial behaviors. The amphibian medial pallium is homologous to the mammalian hippocampus [14,15], though with a more primitive structure that may reflect the simplified architecture of ancestral tetrapods [16,17]. A series of experiments in the toad *Rhinella arenarum* have implicated the medial pallium (in conjunction with other brain regions) in the ability to complete a variety of spatial tasks using visual, geometric, and auditory information [10–13]. However, how these findings translate to real-world navigation in nature has not been explored.

Research across amphibians suggests a wide-spread ability to navigate to sites important for reproduction. Many temperate-region amphibians migrate to breeding sites several kilometers away, such as some salamanders [18–20] and pond breeding toads [21,22]. Some Neotropical poison frogs (family Dendrobatidae) move with high precision within a few hundred meters to transport tadpoles to water pools [23–25]. A variety of sensory modalities facilitate these movements among amphibians, with olfaction [19,21,26] and magnetoreception [27–29] most often implicated. Ability to navigate to home sites outside of the breeding context has also been demonstrated in the cane toad, *Rhinella marina* [30]. However, the sensory modalities used for navigation back to terrestrial home sites, rather than to water bodies related to reproduction, has rarely been investigated.

The cane toad is native to Central and South America and was introduced globally, including in Hawaiʻi and Australia. In Australia, the cane toad can disperse and travel long distances [31,32], sometimes moving 200-meters per night to new locations [33], creating difficulties for species management as the invasive front expands up to 60 km a year [32]. Elsewhere, cane toads rarely move more than 200 meters and often return to the same diurnal shelters [30,34–36]. They do not migrate to breeding ponds like other bufonids in which navigation has been studied [21,22] and their life-histories do not suggest a need for advanced navigation. However, we previously showed that toads in French Guiana are able to return home accurately from a full kilometer, a distance exceeding their routine movements [30]. These abilities and the toad’s extreme prevalence make it an ideal model for studying homing behavior and probing unanswered questions regarding the neural and sensory basis of homing.

In this study, we conduct a field experiment to examine cane toad navigational ability, as well as associated sensory systems and neural activity. We hypothesized that the medial pallium is involved in navigation and predicted that homing toads would have higher neural activity in this brain region compared to non-homing and non-translocated toads. We next hypothesized that olfaction and magnetoreception guide homing behavior, given their function in other bufonids [21,22], and predicted that olfactory or magnetoreception ablations would disrupt the homing ability or accuracy.

## Materials and Methods

### (a) Study sites

Experiments occurred at four sites on the island of O‘ahu in Hawaiʻi. We sampled at two sites on the south of the island (West Loch Golf Course and Hawaiʻi Prince Golf Course) and two sites on the north shore (Waimea Valley and Turtle Bay Golf Course) (Figure S1). Waimea Valley is forested, intercut with paved paths, and has a stream that ran approximately 2-km from a waterfall at the eastern terminus of the field site to a pond at the western terminus with approximately 60-m decline in elevation. All other sites are golf courses with open greens and negligible changes in elevation, interrupted by ponds and forested borders. Turtle Bay was the only site directly bordering the ocean and had wider forested borders than other golf courses. Surface area over which tracking occurred varied between sites (Figure S1). Toads were distributed throughout most of the surface area of each site, suggesting a widely suitable habitat at each location.

We collected daily weather data from the National Oceanic and Atmospheric Administration online database for every day of translocation experiments. The amount of precipitation was significantly different between field sites (Kruskal-Wallis, χ^2^_(3)_= 15.14, p = 0.0017), with the northern sites (Turtle Bay and Waimea Valley) receiving more precipitation than the southern sites. Both northern sites also had lower maximum temperatures (Kruskal-Wallis, χ^2^_(3)_= 39, p = 1.74e-8).

### (b) Animals

Toads were located visually at night (sunset was approximately 19:00 and sunrise approximately 06:30), as cane toads are nocturnal and generally do not exhibit diurnal movements. We photographed the dorsum for re-identification, measured snout-vent-length (SVL) with a tape measure, and measured mass with a portable scale (KUBEI-790, KUBEI, China). Eighty-two toads (40 males, 42 females) were tagged and tracked for at least three days between 18 February and 12 May 2022. Their mean mass was 233.8 g (sd +/− 98.6) and mean SVL was 12.4 cm (+/− 1.6). Toad size was not evenly distributed between sites (Mass: Kruskal-Wallis, χ^2^_(3)_= 19.687, p = 0.0002; SVL: ANOVA, F_(3)_ = 14.67, p = 1.15e-07). Sex was determined by size and male release call and confirmed by gonad identification upon dissection. Females were larger than males (Mass: Wilcoxon, W = 1315.5, p = 1.05e-05; SVL: t-test, t_(77.41)_ = 4.44, p = 2.99e-05). Unless otherwise stated, “size” refers to SVL hereafter.

### (c) Tagging and baseline tracking

Radio tagging and tracking was executed similarly to methodology we previously described [30]. Radio tags (BD2, Holohil Systems Ltd, Carp, ON, Canada) were attached to toads’ waists using silicone tubing and thread (Figure 1A). The tagging process lasted 14.2 min on average (+/− 7.8) after which toads were released at the exact point where they were collected. A GPS point was taken in the ArcGIS Field Maps app (ESRI, Redlands, CA, USA) using a Bad Elf GPS Pro+ gps device (Bad Elf, West Hartford, CT USA) connected by Bluetooth to an iPhone (Apple, Cupertino, CA, USA). Following release, toads were located once in the evening prior to their active period and at least twice each night. Toads were tracked to describe their daily movements and to identify their home areas (termed ‘baseline’ hereafter) between three and 10 days (4.978 +/− 1.48 days) prior to translocation or euthanasia. Any reference to a “baseline point” indicates a coordinate point a toad occupied prior to translocation (i.e. during natural movements). We removed three toads from the analysis of baseline activity because we could not properly observe their movements after they entered burrows.

**Figure 1.**
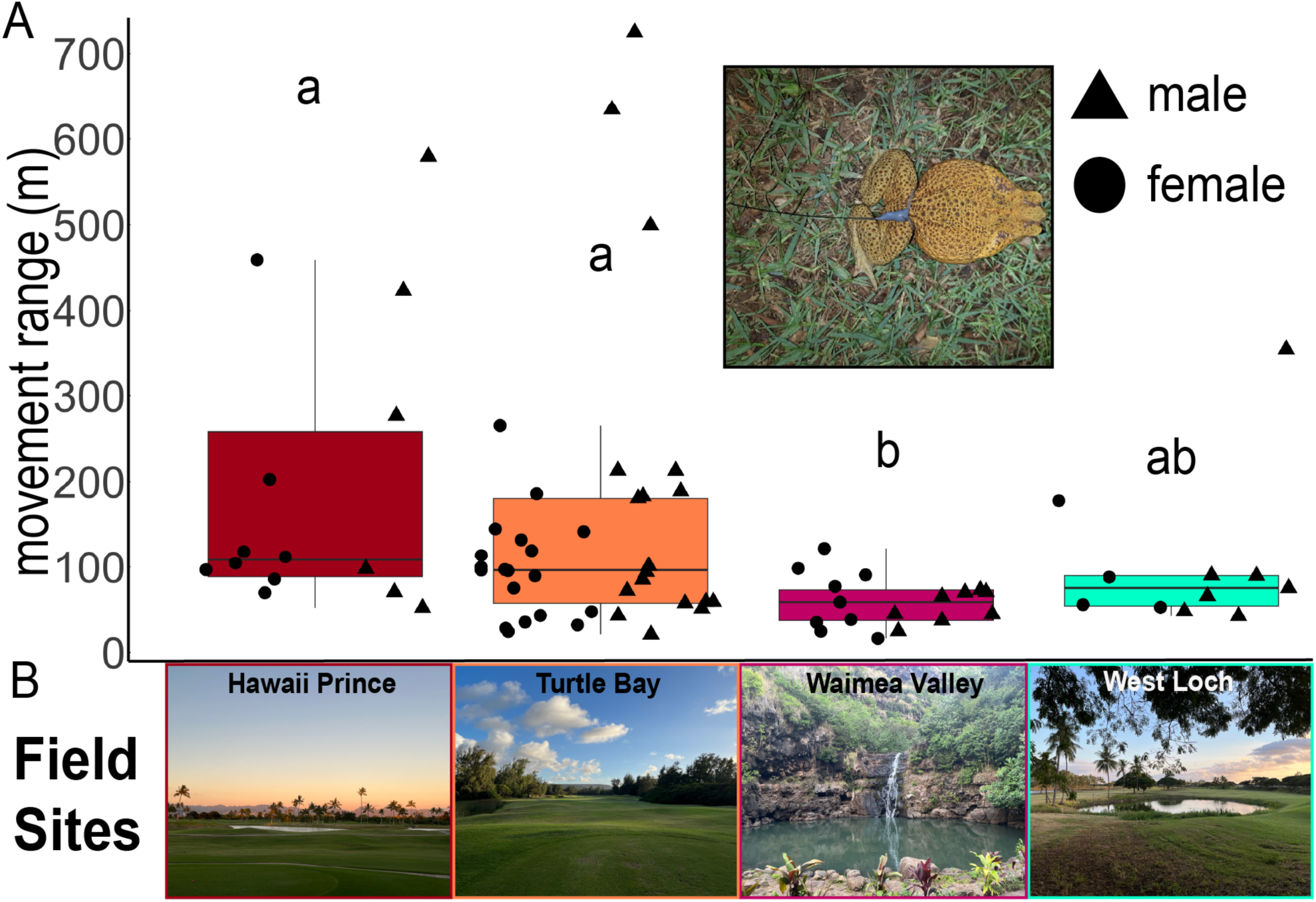
Movement range varies across locations. **(A)** Observed movement range of cane toads at each tracking site. Letters represent significant differences between sites (Kruskal-Wallis, χ^2^_(3)_ = 14.21, p = 0.0026) from post-hoc Dunn’s Test (with Benjamini-Hochberg adjustment). Shapes represent sex (▲ = male, ● = female). Inset shows toad with radio tag. **(B)** Photographs depicting typical landscapes for each site.

### (d) Translocation

Translocations generally occurred between 20:00 and 22:00. Sixty-two toads (32 females and 30 males) were translocated. Toads were caught, their waists examined to ensure no abrasion from the tag belt and placed into an enclosed net cage and walked in an indirect route for 30-60 minutes to a distance approximately 500- or 1000-meters from the point recorded during the baseline tracking period that lay furthest in the direction of translocation. Therefore, the minimum distance to any location of the toad observed prior to translocation matched the translocation distance. Toads gathered in similar locations were translocated in diverging directions as allowed by the geography of each site. A Raleigh’s Test of Uniformity performed on the angles of translocation indicate that they were not preferentially grouped in any direction (p = 0.20). Translocation distances exceed the average movements observed in cane toads in Hawaiʻi [36] and French Guiana [30,35] but are within the range of the maximum movement distances observed, meaning we cannot be certain that all toads were moved to an unfamiliar location. At the release site, the distance from the toad’s nearest baseline point was confirmed on a GPS-connected handheld device, the net cage was left at the release point for 5 minutes, and the toad was gently released from the cage.

Following release, toads were relocated approximately once per hour during nighttime, beginning at 20:00 and continuing until tagged animals had stopped moving (06:40 at the latest). On subsequent days, toads were relocated once in the early evening to determine location before nocturnal movements began. If a toad was not active at night (e.g. not moving or making small, multidirectional movements), the frequency of observations was reduced to every 2-3 hours while tracking moving toads.

Tracking ended once the animal reached a point within 200 meters of one of its recorded baseline positions (homing, n = 33) or 72 hours had passed after translocation (non-homing, n = 27). Homing toads (captured within 200-meters of a baseline point) were euthanized and tissues were collected for downstream analyses (described below) within 30 minutes of crossing the 200-meter threshold. Non-homing toads were sacrificed after 20:00 wherever they were at that time. Two toads in the homing group were unintentionally caught before they crossed the 200-meter threshold (B8 at 233-m and D5 at 218-m) and two more toads were caught and euthanized approximately 300-m from home after encountering physical obstacles following large homeward movements (C8 and F5) but were included in the homing condition when considering behavior and neuronal activity. One toad moved outside of the study area and was euthanized because it became impossible to track, and its behavior and brain data were not included in analysis.

### (e) Sensory Ablations

A subset of translocated toads (n = 43; 22 females and 21 males) received magnetoreception or olfactory ablations to test the roles of these sensory systems in homing. Each modality had experimental (i.e. ablated) animals and control animals receiving sham treatments. Toads were tracked for at least three days prior to the ablation procedure. After the procedure, they were tracked for at least an additional 48 hours before translocation to evaluate any effects of the ablation on space use. Olfactory ablations were performed through zinc sulfate (ZnSO4) treatment, which temporarily damages the olfactory epithelium in amphibians [37,38] and other vertebrates [39,40]. Magnetoreception ablations were carried out by attaching a small Nickel-plated Neodymium magnet (N50, 10 mm x 6 mm x 2 mm, ∼1 g; Super Magnet Man, Pelham, Alabama, USA) with cyanoacrylate based adhesive (EM-150, Starbond, Torrance, CA, USA) to the skin over the skulls of toads, similar to methodology performed in other amphibians [21,41]. Detailed descriptions of ablation and sham procedures can be found in SM1. Toads in ablation experiments were euthanized and tissues collected at the same time points described above.

### (f) Tissue collection

Translocated toads were euthanized at the time points described above. Additional toads were euthanized without translocation after the completion of at least three days of baseline tracking (baseline, n = 12). Toads were euthanized through MS-222 injection (0.7% solution, 300 mg/kg body weight) followed by cervical transection. Brains were immediately dissected, placed in 1 mL of 4% paraformaldehyde (PFA), and kept on ice until they could be placed into a 4℃ refrigerator. Brains were fixed overnight in PFA, washed three times in 1x PBS and then placed into 1x PBS with 0.02% sodium azide, or 1x PBS if transportation to the laboratory site would happen in two days. Brains were kept in 1x PBS with sodium azide < 1 week. Two days prior to travel to the laboratory, the brains were transferred to 1 mL of 30% sucrose dissolved in 1x PBS for cryoprotection. Once the brains sank in sucrose, they were transported on ice to a laboratory at the University of Hawaiʻi at Manoa, where they were embedded in Tissue-Tek OCT Compound (Electron Microscopy Sciences), frozen on dry ice, and stored at −80℃. Seventy-three brains were collected (61 translocated and 12 baseline). One homing toad lost its tag within 200-meters of its home area, so its brain was not collected.

### (g) Immunohistochemistry

Brains were sectioned on a cryostat in 20 µm slices and thaw-mounted onto slides in four series. Slides dried at room temperature for ∼48 hours before being transferred to −80℃ for storage prior to staining. We used immunohistochemistry to detect Phospho-S6 Ribosomal Protein (pS6) as a proxy for neural activity [42] as previously described by our laboratory [43,44] (full protocol presented in SM2). Of the 73 collected brains, two were discarded due to poor tissue quality and four were discarded for being collected at the wrong time point, leaving 67 for analysis.

### (h) Microscopy and Cell Counting

Brains were imaged on a Leica DM6B microscope with brightfield capability at 20x magnification. One hemisphere was selected to image for each brain. Nine sections along the AP axis were chosen based on anatomical landmarks: three from the anterior of the telencephalon (posterior to the olfactory bulb), three from the medial portion, and three from the posterior of the telencephalon (anterior to the ventral hypothalamus) (Figure S2). ImageJ (NIH, Bethesda, MD, USA) was used to count pS6-positive cells in each of six brain regions: medial pallium (Mp), lateral pallium (Lp), dorsal pallium (Dp), medial septum (Ms), lateral septum (Ls), and the striatum (Str) given their role in spatial behavior in toads [10–12] and other vertebrates [45–47]. Delineation of brain regions was based on Northcutt and Kicliter (1980) and Sotelo et al. (2016), which defined pallial regions by Westhoff and Roth (2002) and ventral regions by Moreno and González (2004) [10,15,48,49]. ImageJ was used to calculate the surface area of each region. Densities of pS6-positive neurons for each region were calculated by dividing the number of pS6-positive cells by the area.

For toads in the olfaction ablation experiment, ImageJ was used to count active cells in a 300 µm by 300 µm box positioned at the lateral edge of the granule cell layer in the olfactory bulb (Figure S3) to measure the effect of the olfactory ablation in comparison to the sham ablation.

### (i) Data analysis

#### (i) Toad movement data

GPS points were visualized in ArcGIS online and ArcGIS Pro 2 (ESRI). Spatial and temporal attributes were calculated using the package ‘adehabitatLT’ [50] in R Studio (v 2024.12.0+467, Posit Software, PBC, Sunnyvale, CA) running R (v 4.4.2, R Foundation for Statistical Computing, Vienna, Austria). Several descriptors of toad movement and homing were calculated: cumulative movement, daily movement, movement range, straightness index, homing duration/speed, active homing duration/speed, and similarity of toad’s home range before and after ablation treatment (details of descriptor calculations in SM3). Detailed description of statistical treatment of movement data can be found in SM4.

#### (ii) Cell Count Data

Generalized linear mixed models were used (‘glmmTMB’ in R [51]) to test for differences in brain activity and its relation with behavior, with homing condition, brain region, and their interactions as the main predictors. Post hoc pairwise contrasts (between homing condition groups and ablation treatments) were calculated with estimated marginal means using the “emmeans” package in R [52]. Birds [53] and mammals [54] show variation in hippocampal spatial activity along the anterior-posterior axis, so we also tested for differences in activity along this axis in the Mp. To test if brain activity was related to either straightness or duration of homing, linear mixed effects models were performed testing whether brain activity could be predicted by straightness or duration. Detailed descriptions of statistical treatment of cell count data can be found in SM5.

### (j) Permits and Ethical Statement

All work was done in adherence to Protected Wildlife Permit WL21-16 issued by the State of Hawaiʻi Department of Land and Natural Resources, Division of Forestry and Wildlife. All procedures were approved by the Institutional Animal Care and Use Committee of Stanford University (Protocol #33530).

## Results

### Toads generally exhibit site fidelity

Before performing translocations, we observed baseline movements and found a mean movement range of 129.45 m (s.d. 140.22 m), mean cumulative movement of 547.97 m (s.d. 758.00), and mean daily movement of 114.23 m (s.d. 158.98). Movement range varied between sites (Kruskal-Wallis, χ^2^_(3)_ = 14.21, p = 0.0026), with toads at the Waimea Valley forest site exhibiting smaller ranges than toads at golf courses (Figure 1). There was no sex-based difference in movement range (Wilcoxon, W = 750, p = 0.77). All three movement ranges more than three standard deviations from the mean were male (example in Figure S4). Size did not have an effect on movement range (t_(68.60)_ = −0.814, p = 0.42; Figure S5).

### Toads exhibited long-distance homing

Next, we tested whether toads could navigate back to home areas following displacements of 500- or 1000-m. Out of 62 translocated toads, 34 exhibited homing behavior (Figure 2A-D). Homing success did not vary with sex (χ^2^_(1)_= 0.16, p = 0.69), but did vary with field site (χ^2^_(3)_= 10.33, p = 0.016). Specifically, toads at Hawaii Prince homed with greater success than toads in Turtle Bay (Fisher exact test, p = 0.04) and Waimea Valley (Fisher exact test, p = 0.03). Toads returned home in 3.27 to 77.48 hours (25.50 +/− 22.23) with a mean straightness of 0.81 (+/− 0.16). Considering all toads (including those in ablation experiments), 19 out of 31 toads returned home from 500-meters and 15 out of 30 toads returned from 1000-meters. There was no effect of translocation distance on homing success (Pearson’s Chi-squared, χ^2^_(1)_ = 0.40, p = 0.53). Distance had no effect on homing duration (Wilcoxon, W = 143.5, p = 0.99) (Figure S6A) and toads translocated 1000-meters had faster homing speeds (Wilcoxon, W = 72, p = 0.015; Figure S6B). A linear mixed-effects model accounting for the unequal distribution of toad size across field sites showed no significant effect of size on homing duration (t_(30.52)_ = −0.51, p = 0.61; Figure S7).

**Figure 2.**
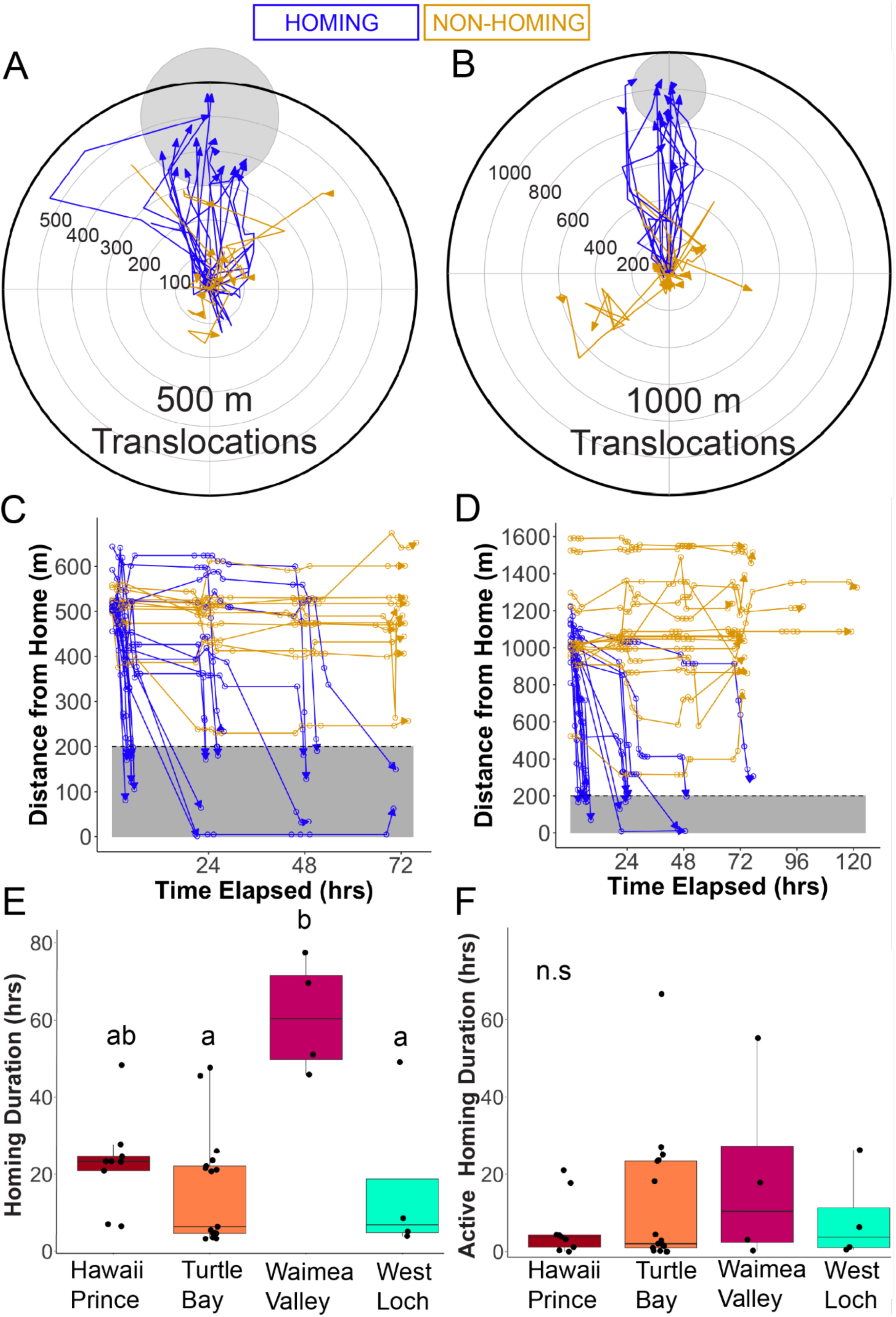
Toads exhibit long-distance homing. Normalized trajectories of toads translocated **(A)** 500-m and **(B)** 1000-m. Plot centers represent translocation release points and the gray circles at the top represent the 200-m buffer threshold around baseline points. 19/31 toads returned from 500-m and 15/30 from 1000-m. Distance to home over time for toads translocated **(C)** 500-m and **(D)** 1000-m. Gray rectangle at the bottom of the plot represents the 200-m buffer threshold around baseline points **(E)** Duration of homing differs between the experimental sites (Kruskal-Wallis, χ^2^_(3)_= 12.38, p = 0.0062) Letters represent significant differences from post-hoc Dunn’s Test (with Benjamini-Hochberg adjustment). **(F)** Active homing time shows no difference between locations (Kruskal-Wallis, χ^2^_(3)_= 0.44, p = 0.93).

Homing speed and duration varied significantly between sites, with toads at Waimea Valley homing slower than toads at Turtle Bay and West Loch (Speed: Kruskal-Wallis, χ^2^_(3)_= 10.52, p = 0.015; Duration: Kruskal-Wallis, χ^2^_(3)_= 12.38, p = 0.0062; Figure 2E). Toads often did not immediately move homeward, so we measured an “active homing speed” using only movement and time after the first 100-meters of homeward movement. Active homing speed and duration did not vary between sites (Speed: Kruskal-Wallis, χ^2^_(3)_ = 4.93 p = 0.18; Duration: Kruskal-Wallis, χ^2^_(3)_= 0.44, p = 0.93), suggesting that the main differences in homing time are due to stationary periods prior to homeward movements (Figure 2F). On average, the time to move at least 100-meters homeward was 64% of the total homing duration.

Twenty-seven of the 62 translocated toads did not exhibit homing behavior. Cumulative movement of non-homing toads averaged 547.07 m (+/− 488.5) and the mean minimum distance from home reached was 719.3 m (+/− 359.5). Mean movement angles of homing toads showed a significant orientation towards the home direction (Rayleigh p = 0.024) and non-homing toads did not (Rayleigh p = 0.83). However, there was only a marginal difference in distribution of angles of movement between homing and non-homing toads (MANOVA, Pillai = 0.092, F_(2,58)_ = 2.95, p = 0.06; Figure S8).

### Olfaction and magnetoreception ablations affect toad movements

We next asked whether ablation of olfaction led to a disruption in spatial behavior and navigation, as has been demonstrated in other amphibians [21,22]. There was no significant difference in movement range pre- and post-ablation for sham (Paired-Wilcoxon, V = 38, p = 0.70) or ablated animals (Paired Wilcoxon V = 36, p = 0.83). However, when comparing the space occupied pre- and post-ablation, ablated toads showed greater divergence from their pre-ablation area than sham animals (Wilcoxon, W = 340, p = 0.021), suggesting that loss of olfaction might disrupt routine movements and space use. There were no significant differences of homing success (χ^2^_(1)_ = 0.73, p = 0.39) or duration (Wilcoxon, W = 12, p = 0.43) between sham and ablated groups (ablated success = 45.5%, sham success = 63.6%)(Figure 3A). After translocation, ablated toads moved more than sham toads when considering homing and non-homing toads together (Wilcoxon, W = 28, p = 0.034) (Figure 3B). Though the sample is too small for statistical comparison, ablated toads that homed showed more cumulative movement (831.90 m +/− 288.07) than sham toads that homed (629.79 m +/− 334.19). Olfactory ablated toads had significantly less pS6-positive cells in their olfactory bulbs than sham toads, confirming reduced olfactory function from chemical ablation (Wilcoxon, W = 93.5, p = 0.0074)(Figure S9).

**Figure 3.**
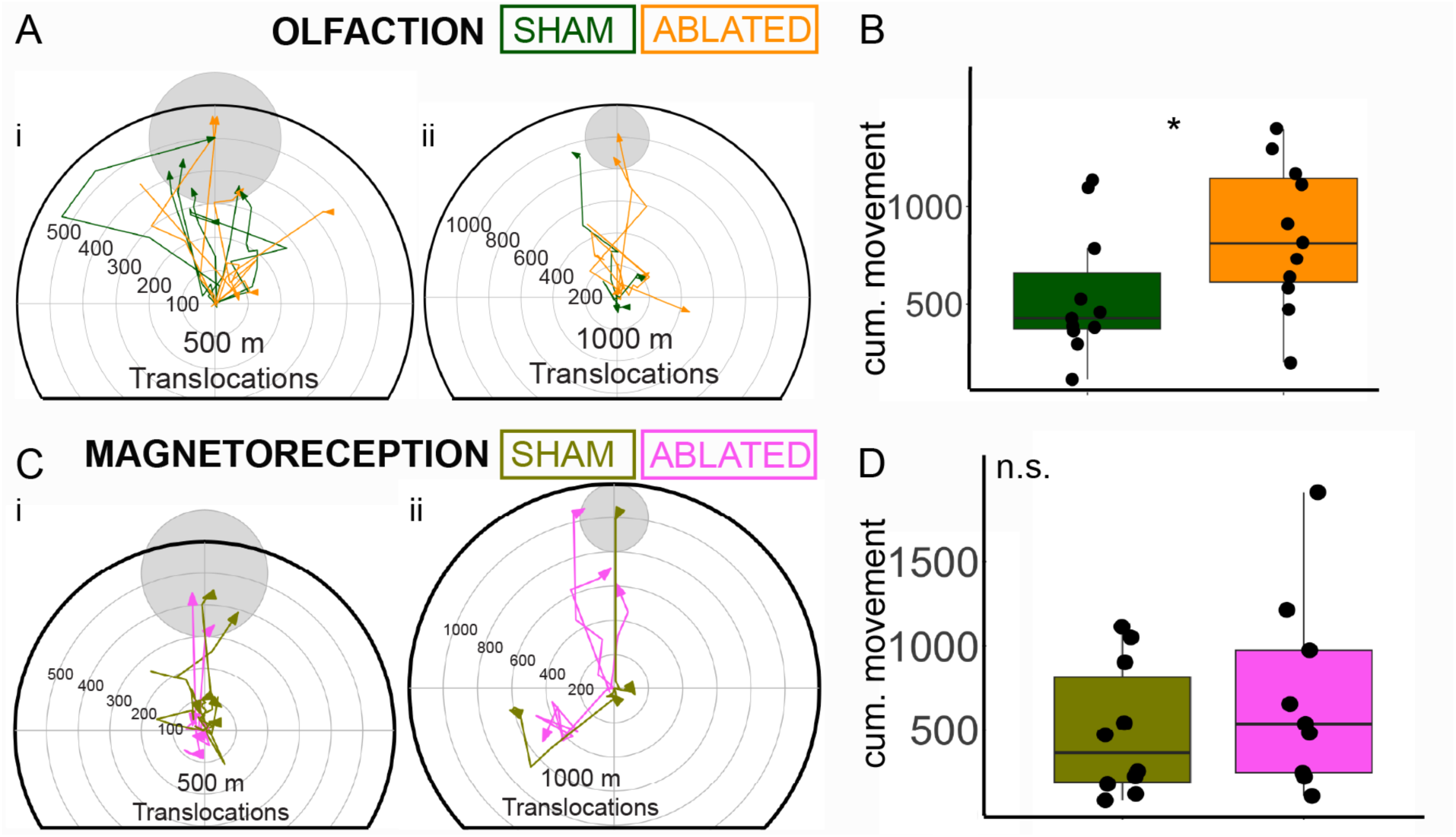
Sensory ablations did not prevent homing. **(A)**Trajectories of toads in the olfactory ablation experiment translocated (i) 500-meters and (ii) 1000-meters. **(B)** Olfaction ablated toads exhibited higher cumulative movement post-translocation than sham toads (Wilcoxon, W = 28, p = 0.034). **(C)** Trajectories of toads in the magnetoreception ablation experiment translocated (i) 500-meters and (ii) 1000-meters. **(D)** Magnetoreception ablated toads exhibited no difference in cumulative movement post-translocation than sham toads (W = 35, p = 0.447).

Magnetoreception is a sensory modality linked to homing in several amphibians [29,55], so we tested the hypothesis that cane toads also use magnetoreception for homing. Movement range was significantly lower in both sham (Paired Wilcoxon, V = 56, p = 0.042) and ablated (Paired Wilcoxon, V = 62, p = 0.0068) animals. There was no significant difference when comparing similarity in positions pre- and post-ablation (Wilcoxon, W = 236, p = 0.90). There were no significant differences of homing success (χ^2^_(1)_ = 1.27, p = 0.26) or duration (Wilcoxon, W = 10, p = 0.57) between sham and ablated groups (ablated success = 55.6%, sham success = 30.0%) (Figure 3C). There was no difference in cumulative movement post-translocation between sham and ablated animals (Wilcoxon, W = 55, p = 0.44)(Figure 3D).

### Pallial and septal brain regions show increased neural activity in homing toads

Given the variability in homing performance, we next asked whether neural activity in brain regions is correlated with homing success by quantifying the number of pS6-positive cells in six brain regions (Figure 4). We first asked whether toads, regardless of sensory treatment, showed differences in brain activity due to homing behavior. We found a significant effect of homing behavior on neural activity in a brain region specific manner (brain region x homing condition: χ^2^_(10)_ = 112.80, p < 2.2e-16) (Table S1). We found that homing animals had more pS6-positive neurons than both non-homing and baseline toads in the medial pallium (Mp), medial septum (Ms), and the lateral septum (Ls) (Table 1). In the lateral pallium (Lp), we also found significant homing-non-homing differences and marginal homing-baseline differences. The dorsal pallium (Dp) and striatum (Str) were not different in any comparison. When we split the Mp into three sections along the A-P axis, we found significant differences between homing toads and the other two conditions for every section, indicating no variation from front to back (Table S2). There were no differences in brain activity across brain regions due to path straightness (Tables S3 and S4). However, the Mp (p = 0.0076) and Ls (p = 0.00076) showed significant differences in activity related to duration of homing compared to other brain regions (Table S5) and duration showed a significant effect of homing on neuronal activity in the Ms (t_(27)_ =-2.55, p = 0.017) and Ls (t_(27)_ = −2.93, p = 0.0069) (Table S6).

**Figure 4.**
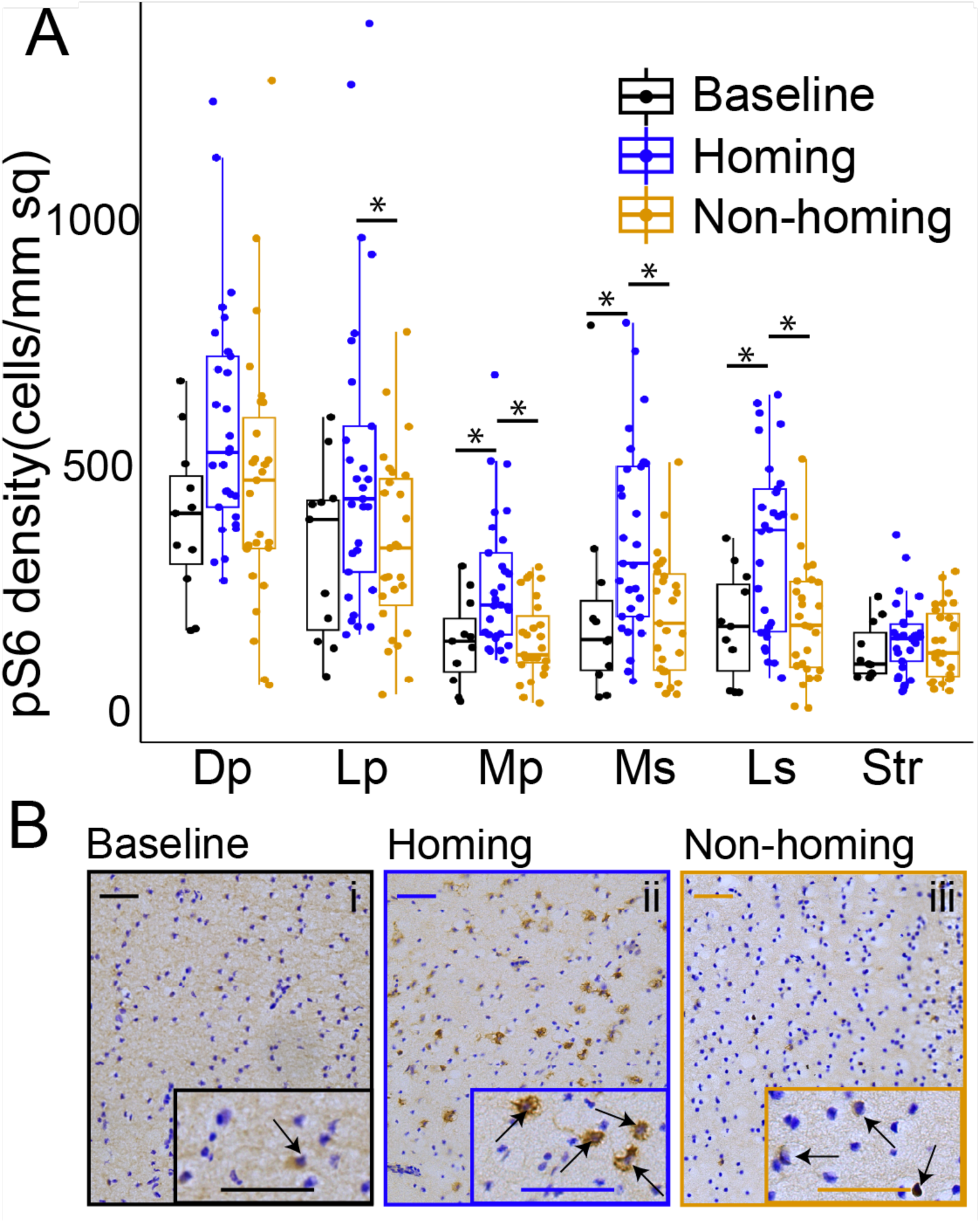
Pallial regions and the septum are more active due to homing behavior. **(A)** pS6-positive cells per mm^2^ averaged per individual for each brain region across homing conditions. Brain activity within brain regions varied significantly due to homing conditions (χ^2^_(10)_ = 112.800, p < 2.2e-16). Stars represent significant pairwise differences within brain regions based on a mixed model with cell count as a response factor. Abbreviations: Dp, dorsal pallium; Lp, lateral pallium; Mp, medial pallium; Ms, medial septum; Ls, lateral septum; Str, striatum. **(B)** Representative examples of stained Mp brain tissue from each homing group. Scale bars represent 50 µm. In inset, arrows indicate pS6-positive cells.

**Table 1.** Pairwise contrasts by Homing Condition in all brains.

| Brain Region | Contrast | estimate | std. error | z-ratio | p-value |
| --- | --- | --- | --- | --- | --- |
| Dp | Baseline-Homing | -0.4071 | 0.202 | -2.013 | 0.1091 |
| Dp | Baseline-Nonhoming | -0.1087 | 0.204 | -0.532 | 0.8558 |
| Dp | Homing-Nonhoming | 0.2984 | 0.153 | 1.957 | 0.1231 |
| Lp | Baseline-Homing | -0.457 | 0.198 | -2.303 | 0.0554 |
| Lp | Baseline-Nonhoming | -0.0521 | 0.201 | -0.26 | 0.9635 |
| Lp | Homing-Nonhoming | 0.405 | 0.15 | 2.703 | 0.0188 |
| Mp | Baseline-Homing | -0.7123 | 0.198 | -3.595 | 0.0009 |
| Mp | Baseline-Nonhoming | -0.096 | 0.2 | -0.48 | 0.881 |
| Mp | Homing-Nonhoming | 0.6163 | 0.15 | 4.12 | 0.0001 |
| Ms | Baseline-Homing | -0.7186 | 0.203 | -3.533 | 0.0012 |
| Ms | Baseline-Nonhoming | -0.058 | 0.206 | -0.282 | 0.9572 |
| Ms | Homing-Nonhoming | 0.6606 | 0.153 | 4.312 | <.0001 |
| Ls | Baseline-Homing | -0.716 | 0.202 | -3.544 | 0.0011 |
| Ls | Baseline-Nonhoming | -0.0598 | 0.204 | -0.293 | 0.9537 |
| Ls | Homing-Nonhoming | 0.6561 | 0.152 | 4.303 | 0.0001 |
| Str | Baseline-Homing | -0.191 | 0.207 | -0.923 | 0.6259 |
| Str | Baseline-Nonhoming | -0.0897 | 0.209 | -0.428 | 0.9038 |
| Str | Homing-Nonhoming | 0.1013 | 0.156 | 0.651 | 0.7917 |

\*Shading represents significant relationships, light shading represents marginal significance

To better account for the effect of sensory ablations on brain activity, we also modeled brain activity in all translocated toads with sensory ablation as a predictor (Table S7). This model predicts significant effects of both homing condition (χ^2^_(5)_ = 119.09, p < 2.2e-16) and ablation (χ^2^_(20)_ = 127.59, p < 2.2e-16) on neural activity in brain region specific manners. However, the magnitude of the effect was insufficient to detect significant pairwise differences in region-specific activity between ablated and sham groups in both olfaction and magnetoreception experiments (Figures S10; Table S8). The model also indicated significant differences between homing and non-homing toads in the Mp (z = 4.138, p < 0.0001), Ms (z = 2.909, p = 0.0036), and Ls (z = 3.916, p = 0.0001), but not the Lp (z = 1.876, p = 0.0607).

## Discussion

In the current study, we examined the neural correlates and sensory basis of cane toad homing. Our results demonstrate that more than half of toads displayed a map-like navigation ability from distances exceeding regular movements. We found that areas associated with navigation in mammals and birds [3,6,45,56] and which have previously been implicated in amphibian lab studies [10–13], such as medial pallium and the septum, show increased activity in homing toads. Contrary to our predictions, we found that olfactory or magnetic ablation did not disrupt homing, suggesting this behavior is multimodal, as has been demonstrated in other amphibians [21,57]. These results contribute to filling in gaps in the understanding of amphibian spatial behaviors and neural architecture of vertebrate spatial cognition.

### (a) Homing in toads

We found that most toads could navigate home from up to 1 kilometer. Amphibian homing abilities have previously been linked to the spatial challenges of parental care and seasonal migrations [23,57]. However, cane toad homing suggests that refined navigation may not be reserved for species with specialized reproductive behaviors but is instead more widely present across amphibians. Cane toads outside the Australian invasive front typically moved less than 200-meters, but rare movements of 500-meters to 2-kilometers have been observed [30,35,36]. Thus, our translocation distances moved toads beyond the area of regular movements, but were still within the range that toads could occasionally explore. In addition, toads were found distributed throughout each field site, rather than clustered at specific sites. Therefore, for most translocated toads, suitable habitat was available closer to the release site than home, which suggests that toads were navigating to specific locations and not simply finding the nearest suitable habitat.

Many toads did not home, suggesting strong variability in ability or motivation to move back home. Some variation may be attributable to environmental features, as the more wooded sites (Waimea Valley and Turtle Bay) had lower levels of success than Hawaii Prince. Vegetation may create landscape resistance and occlude sensory information or provide additional shelter that reduces motivation to return. This is in line with the results of a homing experiment with *R. marina* in primary rainforest [30], where homing was generally slower than that observed in this experiment. Turtle Bay and Waimea Valley also experienced more precipitation than West Loch and Hawaii Prince. More available water could reduce pressure to return to previously occupied areas, resulting in lower rates of homing (in Australia, availability of water has been shown to influence daily movements of cane toads [58]). However, individual variation in ability, experience, and motivation leading to variable homing success might also play in an important role. Inter-individual variability unaccounted for by environmental variation has been often observed in amphibian navigation studies [23,59] and is likely caused by individual differences that should be further explored.

### (b) Use of sensory systems in homing

Olfaction and magnetoreception have been implicated as important senses in navigation studies in many amphibians [19–22,27,29,57] and other vertebrates [60–64]. Ablation experiments strongly suggested that *B. bufo* relies on these senses to orient towards breeding ponds [21], while *B. japonicus* seemed to rely solely on olfaction [22]. In a variety of salamander species, both magnetoreception and olfaction are used to orient and return to breeding streams [19,20,27,28]. However, in our study, these ablations did not disrupt *R. marina* homing. If either is used in cane toad navigation, they likely work in combination with other senses or can compensate for each other.

Olfaction is used in multiple ways in navigation. Birds use odors distributed by wind to determine goal locations [60]. Mammals sample odor information through sniffing to gain information about their environment [61]. For amphibians, cues can come directly from aquatic navigation goals, as seems to be the case in other toad species [21,22] and poison frogs [26]. Some salamanders may smell pheromones left by themselves or conspecifics [19]. Although we did not observe a reduced success in homing with olfactory ablations, ablated toads showed less similarity in space occupied post-ablation and moved more post-translocation than sham toads. These results suggest that olfactory cues may help toads move in more direct paths and are important for recognizing home areas. This is somewhat different from what has been observed in migratory bufonids: when olfaction was ablated, *B. bufo* could not orient correctly but still moved in a direct path [21] and *B. japonicus* was completely disoriented [22]. Differences in the specific utility of olfaction during navigation may arise from the observed variations in life history.

Magnetic orientation has been demonstrated in several amphibians [27,65,66], including other bufonids [21,67], but we found no evidence that magnetic field disruption impacts long-distance homing in cane toads. Other vertebrates, such as pigeons, sea turtles, and cetaceans exhibit long range navigation using magnetic cues [62–64]. Much of the functional understanding of magnetoreception comes from salamanders [27,29,65], which can use magnetoreception as both compass and map senses (similarly to sea turtles and birds), even though their movements are on much smaller scales [55,68]. The effect of magnetoreception ablation on homing in other toad species is variable: *B. bufo* showed less accuracy in orientation following translocation [21], while *B. japonicus* showed no response to ablation [22]. We cannot discount magnetoreception in *R. marina* on the basis of the current experiment, as the sham group performed poorly with only 30% of toads returning home. Although we used methodology previously deployed in other amphibians [21,41], both ablated and sham groups moved less following ablation. Treatment may have had an unintended consequence on behavior, possibly through physically covering the pineal gland, which contains photoreceptors [69] that may play a role in light dependent magnetic compass orientation [28]. Future experiments are necessary to further explore the possible role of the pineal gland in navigation or determine other factors affecting locomotion and navigation.

Other senses may play a role in homing behavior. For example, both male and female cane toads exhibit phonotaxis to male calls [12,70]. However, we observed toads distributed over several ponds and inconsistent calling, making the use of auditory cues unlikely. Vision may also play a role and is used by toads and poison frogs in laboratory arenas, both in the context of identifying feature cues and environmental geometry [10,71,72]. It is unlikely to be the primary sense involved in homing, as the visual information toads receive (ground-level, at night) is very limited in relation to the scale of the translocation distance. While environmental geometry could play some role in cane toad navigation, geometry in amphibians and rodents is considered on much smaller scales in enclosed arenas [10,72,73] and does not directly translate to large scale movements in complex environments. In addition, migratory amphibians with visual ablations are able to return home [20,21]. However, vision may be used to sense celestial cues, which provide compass information to some amphibians [57,74]. Further study is necessary to assess the functionality of sensory modalities during navigation.

### (c) Conserved role of pallial brain regions in spatial tasks

Our study contributes to the accumulating evidence that medial pallium is functionally homologous to the mammalian hippocampus. The hippocampus performs many navigation-related functions, such as encoding place [1,45] and consolidating spatial memory [2]. In pigeons, hippocampal lesions impair homing [5] fish and turtles require homologous regions to solve a spatial task [9]. Extensive neuronal activation and lesion studies in the toad *Rhinella arenarum* suggest that the Mp is used in spatial tasks involving environmental geometry [10], feature cues [11], and conspecific mating calls [12]. The Mp is also shown to have increased activity in tadpole-transporting poison frogs, a parental care task that requires spatial memory [44]. Our work corroborates this accumulating evidence that the Mp is involved in the regulation of spatial behaviors in amphibians.

In our experiment, the lateral pallium showed increased activity in homing animals, while the dorsal pallium did not. There is ongoing uncertainty regarding specific functions and homologies of pallial divisions in amphibians [15,75]. Some evidence points to the Dp being homologous to the mammalian entorhinal cortex and subiculum [76], though it has also been considered similar to the general cortex [75,77]. Meanwhile, the Lp is proposed to be homologous to the piriform or olfactory cortex [14,75,77]. However, we saw no evidence of activity changes in the Lp associated with olfactory ablations (Figure S10, Table S8). Sotelo et al. 2024 [13] presents evidence from the toad *R. arenarum* [10] and the newt *Triturus alpestris* [78] suggesting the Dp and Lp are necessary for learning associated with visual stimuli and thus support the Mp in navigation. The potential homologies these regions represent to the mammalian brain and their interconnectedness with the Mp [14] further support the idea that they may be involved in spatial functions. We found that lateral, but not dorsal, pallium was associated with long-distance homing, suggesting distinct and non-visual functions of these regions in relation to navigation.

Our results showed increased activity in the septum in homing animals and increasing activity with faster homing. Both medial and lateral portions of the septum have been implicated in spatial tasks in rodents, with the Ms integrating circuitry important to path integration [56] and the Ls exhibiting reward-related place fields [79]. The septum has also been implicated in navigation in amphibians: in *R. arenarum,* the Ms, but not Ls, showed heightened activity when animals located goal locations in relation to mating calls [12], environmental geometry, and feature cues [13]. Our results differ in this regard, as we find that both the Ms and Ls show increased activity in homing toads compared to non-homing and non-translocated toads. These data corroborate findings in *R. arenarum* but also suggest that the Ls may govern navigation that requires large movements and not necessarily smaller scale spatial learning.

One important caveat of our experiment is the relationship of the behavior and the timing of the endpoint. Homing animals were often stationary before moving back in a straight line, suggesting that the animals gain their bearings during this time. The snapshot of brain activity captured by our pS6 immunohistochemistry does not account for this period, but for brain activity happening during active homing. In other animals, the hippocampal formation is responsible for specific homing processes. For example, in homing pigeons, it governs use of landmarks when approaching home, but not initial homeward orientation [6]. In future toad experiments, it would be useful to compare active-homing brain activity to that of the pre-homing period to identify a neural signature of orientation.

## Conclusion

Our work combines intensive field-based telemetry with laboratory analysis of brain activity to study the neural and sensory basis of navigation. We demonstrate that cane toads (a) are capable of long-distance navigation to specific sites, (b) exhibit conserved function of the pallial and septal regions in relation to spatial behavior, and (c) likely have a multimodal basis of navigation. In the future, lesion experiments are needed to identify the specific function of pallium brain regions in navigation. In addition, further lab-based studies incorporating electrophysiology may characterize how neurons encode spatial information in amphibians. The current work provides a foundation upon which to begin further examinations of the neural and sensory basis of amphibian navigation through the integration of field and laboratory data.

## Supporting information

R Scripts for Analysis

Datasheets

## Acknowledgements

The authors respectfully and with reflection offer a Hōʻoia ʻĀina, a Land Acknowledgement, acknowledging that this work was performed in Hawaiʻi, an indigenous space whose original people are today identified as Native Hawaiʻians. We also acknowledge that the laboratory portion of this research was conducted on the ancestral lands of the Muwekma Ohlone people at Stanford. We understand the implications of the historical and present colonialism the Ohlone people experience and celebrate their continued stewardship of their lands. We thank Dr. Amber Wright at the University of Hawaiʻi at Mānoa for providing us with lab spaces and community. We thank the representatives of our field sites who made our work possible: George Crisologo, William Suckoll, Travis Joeger, and Chad Middleton. and all employees who assisted us. We also thank Camilo Rodríguez Lopez for his counseling on statistical matters and Neil Khosla for reading preliminary drafts of this work.

## Data Availability

All data and code have been uploaded as supplemental information.

## Author contributions

Conceptualization: D.A.S., A.P., L.A.O; Methodology: D.A.S.; Formal analysis: D.A.S.; Investigation: D.A.S., D.P.M., C.G.; Resources: L.A.O.; Data curation: D.A.S., D.P.M., C.G; Writing – original draft: D.A.S.; Writing – review and editing: D.A.S., L.A.O., A.P., D.P.M., C.G.; Visualization: D.A.S.; Supervision: L.A.O.; Project administration: L.A.O.; Funding acquisition: D.A.S., L.A.O.

