## Supplementary material for "Neural and sensory basis of homing behavior in the invasive cane toad, *Rhinella marina*": R Scripts for Analysis: Master Script- Neural and sensory basis of homing behavior in the invasive cane toad, Rhinella marina.R

```
#R Script for Neural and sensory basis of homing behavior in the invasive cane toad, Rhinella marina
rm(list=ls()) 
setwd("~/Documents/DocumentsDanielWork/HI_2022")  

#Load required packages
library(devtools) 
library(tidyr)
library(ggplot2)
library(DHARMa)
library(emmeans)
library(psych)
library(lme4)
library(nlme)
library (effects)
library (cowplot)
library(interactions)
library(ggpubr)
library(gtsummary)
library(mblm)
library(rcompanion)
library(coda)
library(rjags)
library(BayesFactor)
library(Bolstad)
library(dplyr)
library (trajr)
library(adehabitatLT)
library(adehabitatHR)
library(raster)
library(OneR)
library(chron)
library(datetime)
library(knitr)
library(spdep)
library(lubridate)
library(ggforce)
library(sp)
library(sf)
library(geosphere)
library(glmmTMB)
library(ggsignif)
library(car)
library(MASS)
library(broom)
library(lmerTest)
library(circular)
library(FSA)

#Set colors
loc_colors <- c(
  "West Loch" = "aquamarine",
  "Hawaii Prince" = "brown",
  "Waimea Valley" = "#C03A76FF",
  "Turtle Bay" = "#FD9A6AFF"
)

ab_colors <- c(
  "None" = "black",
  "MAE" = "violet",
  "MAC" = "yellow4",
  "MAE*" = "darkgray",
  "OAE" = "orange",
  "OAC" = "darkgreen"
)

con_colors <- c(
  "Baseline" = "black",
  "Homing" = "blue",
  "Lost" = "goldenrod"
)
####################################################################################################
#PART 1
#Analysis of translocation related spatial activity
###################################################################################################
#read the CSVs containig all the points
toad_info <- read.csv("~/Documents/DocumentsDanielWork/HI_2022/ArcGIS Exports/TransToadInfo.csv")

toad_info_df <- as_tibble(toad_info) %>%
  dplyr::select(id = id, sex = Sex, mass = Mass, svl = SVL, loc = Location, homing = Homing, con = Condition, pair = Pair) %>%
  arrange(id) %>%
  group_by(id)

#Toad B7 moved off the golf course and was not considered for the full amount of time
toad_info_df <- toad_info_df[c(-match("B7", toad_info_df$id)),]
table(toad_info$Sex)
###########################################################################################################
#Load Related Datasets

#Information on collected brains
brains2 <- read.csv("~/Documents/DocumentsDanielWork/HI_2022/Brain Info/Brains.csv")
brains2$id <- brains2$Toad
brains2$Toad <- NULL

#Spatial information from baseline tracking
base_summary <-read.csv("~/Documents/DocumentsDanielWork/HI_2022/base_summary.csv")

#join brain info with other toad characteristics
toad_info_df <- left_join(toad_info_df, brains2)

#Fill in missing data for B3 (brain not collected)
z <- match("B3", toad_info_df$id)
toad_info_df$homing[z] <- 1
toad_info_df$Distance[z] <- 1000
toad_info_df$Condition[z] <- "Homing"
toad_info_df$Ablation[z] <- "None"
###########################################################################################################
#Load translocation coordinate data
Translocation_coords <- read.csv("~/Documents/DocumentsDanielWork/HI_2022/ArcGIS Exports/Translocation_Coordinates.csv")

#clean dataframe of useless variables
toads_all <- Translocation_coords[,c("ID", "Timestamp_E", "POINT_X", "POINT_Y", "Grp")]
names(toads_all )[names(toads_all ) == 'ID'] <- 'id'

#Dataframes with coordinates to calculate the angle animal must travel from translocation point to home
home_angle_all<- read.csv("~/Documents/DocumentsDanielWork/HI_2022/ArcGIS Exports/Transect_Correction.csv")

#convert the timestamp to a POSIXct object so it can be used in adehabitatlt
da <- as.POSIXct(strptime(toads_all$Timestamp_E,"%Y-%m-%d %H:%M"))
da_ha <- as.POSIXct(strptime(home_angle_all$Timestamp,"%Y-%m-%d %H:%M"))

#create a trajectory item for each toad
traj <- as.ltraj(xy = toads_all[,c("POINT_Y","POINT_X")], date = da, id = toads_all$id)   

#create trajectories to determine home angle
home_angle_traj <- as.ltraj(xy=home_angle_all[,c("y","x")], date = da_ha, id = home_angle_all$ID)

######################################################################################################
#create summaries and calculate the total duration observed during tracking
traj_all_summary <- summary(traj)
traj_all_summary$traj.time <- traj_all_summary$date.end - traj_all_summary$date.begin

traj_all_summary <- left_join(traj_all_summary, toad_info_df)

######################################################################################################
#convert the ltraj object back to data.frame with all trajectory attributes
traj_df <- ld(traj)
home_ang_df <- ld(home_angle_traj)
######################################################################################################
#clean up the dataframes from NAs and unneeded variables
toads_df <- as_tibble(toads_all) %>%
  dplyr::select(id = id, dt = Timestamp_E, x_utm = POINT_X, y_utm = POINT_Y) %>%
  arrange(id, dt) %>%
  group_by(id)

home_ang_df %>%
  filter(!is.na(abs.angle))%>%
  dplyr::select(id, abs.angle, dist) -> home_ang__df2

#combine traj_all_summary with home angle and make the angles all positive
traj_all_summary <- left_join(traj_all_summary, home_ang__df2)
traj_all_summary %>%
  mutate(trans_angle = ifelse(abs.angle > 0, abs.angle, 2*pi - abs(abs.angle))) -> traj_all_summary 

#check for distribution of translocation angles
traj_all_summary$trans_angle <- circular(traj_all_summary$trans_angle, type = "angles", units = "radians")
plot(traj_all_summary$trans_angle, bins = 30, main = "Circular Histogram of Angles")
rayleigh_test <- rayleigh.test(traj_all_summary$trans_angle)
print(rayleigh_test)

plot(traj_all_summary[traj_all_summary$loc == "West Loch",]$trans_angle, bins = 30, main = "Circular Histogram of Angles")
rayleigh_test_wl <- rayleigh.test(traj_all_summary[traj_all_summary$loc == "West Loch",]$trans_angle)
print(rayleigh_test_wl)

plot(traj_all_summary[traj_all_summary$loc == "Waimea Valley",]$trans_angle, bins = 30, main = "Circular Histogram of Angles")
rayleigh_test_wv <- rayleigh.test(traj_all_summary[traj_all_summary$loc == "Waimea Valley",]$trans_angle)
print(rayleigh_test_wv)

plot(traj_all_summary[traj_all_summary$loc == "Hawaii Prince",]$trans_angle, bins = 30, main = "Circular Histogram of Angles")
rayleigh_test_hp <- rayleigh.test(traj_all_summary[traj_all_summary$loc == "Hawaii Prince",]$trans_angle)
print(rayleigh_test_hp)

plot(traj_all_summary[traj_all_summary$loc == "Turtle Bay",]$trans_angle, bins = 30, main = "Circular Histogram of Angles")
rayleigh_test_tb <- rayleigh.test(traj_all_summary[traj_all_summary$loc == "Turtle Bay",]$trans_angle)
print(rayleigh_test_tb)

#circular regression to test whether success related to direction
traj_all_summary$loc <- as.factor(traj_all_summary$loc)
model <- glm(homing ~ trans_angle + loc, family = binomial(link = "logit"), data = traj_all_summary)
summary(model)

#join  with toad info
toads_df <- left_join(toads_df, toad_info_df, by = "id")
#####################################################################
#calculate cumulative distance traveled for each toad
for (i in sequence(length(traj_all_summary$id))){
  traj_sub <- traj[which(id(traj) == traj_all_summary$id[i])]
  traj_all_summary$cumulative[i] = sum(na.omit(traj_sub[[1]]$dist))
}

#################################################################################
#calculate displacement transect length from release pt to return pt
for (i in sequence(length(traj_all_summary$id))){
  traj_sub <- traj[which(id(traj) == traj_all_summary$id[i])]
  x <- traj_sub[[1]]$x
  y <- traj_sub[[1]]$y
  lngt = length(traj_sub[[1]]$x)
  traj_all_summary$return_transect[i] = pointDistance(c(x[1],y[1]),c(x[lngt],y[lngt]),lonlat = FALSE, allpairs =FALSE)
}

######################################################################################################
#calculate straightness of path
traj_all_summary$straightness <- (traj_all_summary$return_transect) / (traj_all_summary$cumulative)
######################################################################################################
#calculate time elapsed, and distance from home at every time
toads_df <- toads_df %>%
  group_by(id) %>%
  mutate(time_elapsed = as.numeric(difftime(dt, dt[1], units =  "hours")))

#filter out coordinates of nearest baseline pt ("home")
home_pts <- ld(home_angle_traj)

home_pts %>%
  filter(is.na(abs.angle))%>%
  dplyr::select(id, home_x = x, home_y = y) -> home_pts

home_dist <- c(rep(0, length(toads_df$id)))
i <- 1
for (q in sequence(length(traj))){
  for (r in sequence(length(traj[[q]]$dist))){
    home_dist[i] <- pointDistance(c(traj[[q]]$x[r],traj[[q]]$y[r]),c(home_pts$home_x[q],home_pts$home_y[q]),lonlat = FALSE, allpairs =TRUE)
    i <- i+1
  }}
toads_df$home_dist <- home_dist
###############################################################################
#pick out the coordinates of home_pts and join to the dataframes
toads_df <- right_join(toads_df, home_pts)
###############################################################################
# calculate duration of homing and add it to summary tables- this takes into account the first time the toads 
#get within 200 m of home
for (i in sequence(length(traj_all_summary$id)))
{
  sub <- toads_df[toads_df$id == traj_all_summary$id[i],]
  sub <- sub[sub$home_dist < 200, ]
  traj_all_summary$duration[i] <- sub$time_elapsed[1]
}

#Counting C8 as homing-  reached ~300 m and sacced to preseve brain activity
traj_all_summary[traj_all_summary$id == "C8",]$duration <- 77.483333 

#F5 counted as homing- was beginning to move laterally to go around a big pond ~250 m away
traj_all_summary[traj_all_summary$id == "F5",]$duration <- 6.45

#B8- unintentionally grabbed it at 230 meters- still count as homing brain
#D5- unintentionally grabbed early
traj_all_summary[traj_all_summary$id == "B8",]$duration<- 27.8166667
traj_all_summary[traj_all_summary$id == "D5",]$duration <- 23.583333

#Set duration as 80 hours for all toads that didn't home
for (i in sequence(length(traj_all_summary$id))){
  if (traj_all_summary$homing[i] == 0){
    traj_all_summary$duration[i] <- 80
  }
}

#######################################################################################################################################
#calculate speed of homeward movement for all homing toads
for (i in sequence(length(traj_all_summary$id))){
  if (traj_all_summary$homing[i] == 1){
    traj_all_summary$homing_speed[i] <- (traj_all_summary$return_transect[i]/traj_all_summary$duration[i])
  }
  else{
    traj_all_summary$homing_speed[i] <- NA
  }
}
#######################################################################################################################################
#calculate minimum distance from home reached for all toads
toads_df <- toads_df %>% arrange(id, time_elapsed)

min_dist <- toads_df %>%
  group_by(id) %>%
  summarise(min_dist = min(home_dist))

traj_all_summary <- left_join(traj_all_summary,min_dist)
#######################################################################################################################################
#Calculating time to move 100 m homeward

# Sort by "id" and "time_elapsed" to ensure data ordered correctly
toads_df <- toads_df %>% arrange(id, time_elapsed)

# Create new column to store first "home_dist" value for each toad (initial distance from home)
toads_df <- toads_df %>%
  group_by(id) %>%
  mutate(initial_home_dist = first(home_dist))

# Find point when  toad has first moved 100 meters closer to home
result <- toads_df %>%
  filter(home_dist <= initial_home_dist - 100) %>%
  group_by(id) %>%
  summarise(dist_to_home = first(initial_home_dist), time_to_100 = min(time_elapsed), dist_post_100 = first(home_dist), ts100 = first(dt))%>%
  mutate(act_100 = dist_to_home - dist_post_100)

traj_all_summary <- left_join(traj_all_summary,result)

#Calculate cumulative distance pre 100 meter homeward movement
traj_all_summary$cum_pre_100 <- NA

for (i in seq(length(traj_all_summary$id))){
  sub <- traj_df %>%
    filter(id == traj_all_summary$id[i])
  
  if(!is.na(traj_all_summary$ts100[i])){
    j <- which(sub$date == traj_all_summary$ts100[i])
    traj_all_summary$cum_pre_100[i] <- sum(na.omit(sub$dist[1:j-1]))
  }
}

#proportion of time of total trajectory that 100 meter movement makes up
traj_all_summary$prop100 <- traj_all_summary$time_to_100/traj_all_summary$duration
summary(traj_all_summary[traj_all_summary$homing==1,]$prop100)
sd(traj_all_summary[traj_all_summary$homing==1,]$prop100)
#######################################################################################################################################

#CALCULATE THE ACTIVE HOMING TIME
traj_all_summary$homing_time <- traj_all_summary$traj.time - traj_all_summary$time_to_100 
traj_all_summary$homing_dist <- traj_all_summary$return_transect - traj_all_summary$act_100 
traj_all_summary$homing_speed_post100 <- as.numeric(traj_all_summary$homing_dist)/as.numeric(traj_all_summary$homing_time)
for (i in seq(length(traj_all_summary$id))){
  if (is.infinite(traj_all_summary$homing_speed_post100[i])){
    traj_all_summary$homing_speed_post100[i] <- traj_all_summary$homing_speed[i]
  }}

#######################################################################################################################################
#subsets for downstream reporting
homing <- traj_all_summary[traj_all_summary$homing == 1,]
non_homing <- traj_all_summary[traj_all_summary$homing == 0,] 
homing500 <- homing[homing$Distance ==500,]
homing1000 <- homing[homing$Distance ==1000,]

###############################################################################
###TESTS
min(homing$traj.time)
max(homing$traj.time)
summary(as.numeric(homing$traj.time))
sd(homing$traj.time)
#no outliers for traj.time
Min_time <- mean(as.numeric(homing$traj.time)) - (3*sd(as.numeric(homing$traj.time)))
Max_time <-  mean(as.numeric(homing$traj.time)) + (3*sd(as.numeric(homing$traj.time)))


traj_all_summary$loc <- as.factor(traj_all_summary$loc)
traj_all_summary$homing <- as.factor(traj_all_summary$homing)

#significant
table_data <- table(traj_all_summary$loc, traj_all_summary$homing)
a <- chisq.test(table_data)
a
pairwiseNominalIndependence(table_data, fisher = TRUE, chisq = TRUE, method = "bonferroni")

table_data2 <- table(traj_all_summary$sex, traj_all_summary$homing)
b <- chisq.test(table_data2)
b
pairwiseNominalIndependence(table_data2, fisher = TRUE, chisq = TRUE, method = "bonferroni")

summary(as.numeric(homing$straightness))
sd(as.numeric(homing$straightness))
#no outliers for straightness
Min_straight <- mean(homing$straightness) - (3*sd(homing$straightness))
Max_straight <-  mean(homing$straightness) + (3*sd(homing$straightness))

table(traj_all_summary[traj_all_summary$Distance == 500,]$homing)
table(traj_all_summary[traj_all_summary$Distance == 1000,]$homing)

chisq.test(traj_all_summary$Distance, traj_all_summary$homing)

#no speed outliers
Min_speed <- mean(as.numeric(homing$homing_speed)) - (3*sd(as.numeric(homing$homing_speed)))
Max_speed<-  mean(as.numeric(homing$homing_speed)) + (3*sd(as.numeric(homing$homing_speed)))
#compare speed by location
kruskal.test(homing_speed~loc, data = homing)
dunnTest(homing_speed~loc, data = homing, method  = "bh")
wilcox.test(homing_speed~Distance, data = homing, exact = FALSE)

#no duration outliers
Min_dur <- mean(as.numeric(homing$duration)) - (3*sd(as.numeric(homing$duration)))
Max_dur<-  mean(as.numeric(homing$duration)) + (3*sd(as.numeric(homing$duration)))
summary(homing$duration)
#compare duration by location
kruskal.test(duration~loc, data = homing)
dunnTest(duration~loc, data = homing, method  = "bh")
wilcox.test(duration~Distance, data = homing, exact = FALSE)

#model svl effect on duration
size_duration1 <- lm(duration ~ svl, data = homing)
summary(size_duration1)
#mixed model accounting for site
size_duration2 <- lmer(duration ~ svl + (1|loc), data = homing)
summary(size_duration2)

#one active homing speed outlier
Min_actspeed <- mean(as.numeric(homing$homing_speed_post100)) - (3*sd(as.numeric(homing$homing_speed_post100)))
Max_actspeed<-  mean(as.numeric(homing$homing_speed_post100)) + (3*sd(as.numeric(homing$homing_speed_post100)))
summary(as.numeric(homing$homing_speed_post100))
#compare active homing speed by location with outliers removed
kruskal.test(homing_speed_post100~loc, data = homing[homing$homing_speed_post100 < Max_actspeed,])
#wuthout outliers removed
kruskal.test(homing_speed_post100~loc, data = homing)

#one active homing time outlier
Min_actsdur <- mean(as.numeric(homing$homing_time)) - (3*sd(as.numeric(homing$homing_time)))
Max_actdu<-  mean(as.numeric(homing$homing_time)) + (3*sd(as.numeric(homing$homing_time)))
summary(as.numeric(homing$homing_time))
#compare active homing time by location
kruskal.test(homing_time~loc, data = homing[homing$homing_time < Max_actdu,])
kruskal.test(homing_time~loc, data = homing)

#one active homing speed outlier
summary(homing$homing_speed_post100)
wilcox.test(homing_speed_post100~Distance, data = homing[homing$homing_speed_post100 < Max_actspeed,], exact = FALSE)
wilcox.test(homing_speed_post100~Distance, data = homing, exact = FALSE)

#no outliers time to 100 m
Min_timeto100 <- mean(as.numeric(homing$time_to_100)) - (3*sd(as.numeric(homing$time_to_100)))
Max_timeto100<-  mean(as.numeric(homing$time_to_100)) + (3*sd(as.numeric(homing$time_to_100)))
summary(homing$time_to_100)
wilcox.test(time_to_100~Distance, data = homing, exact = FALSE)

###############################################################################
#looking at non-homing toads
summary(as.numeric(non_homing$cumulative))
sd(non_homing$cumulative)

#no outliers, min_dist
summary(non_homing$min_dist)
sd(non_homing$min_dist)
Min_mindist <- mean(as.numeric(non_homing$min_dist)) - (3*sd(as.numeric(non_homing$min_dist)))
Max_mindist<-  mean(as.numeric(non_homing$min_dist)) + (3*sd(as.numeric(non_homing$min_dist)))

kruskal.test(min_dist~loc, non_homing)#ns

###############################################################################
#Look at magnet and olfaction toads
traj_MA_summary <- traj_all_summary[traj_all_summary$Ablation == "MAC"|traj_all_summary$Ablation == "MAE",]
traj_OA_summary <- traj_all_summary[traj_all_summary$Ablation == "OAC"|traj_all_summary$Ablation == "OAE",]

chisq.test(traj_OA_summary$Ablation, traj_OA_summary$homing, correct=FALSE)
chisq.test(traj_MA_summary$Ablation, traj_MA_summary$homing, correct=FALSE)

#no outliers, cumulative movement OA
Min_OAcum <- mean(as.numeric(traj_OA_summary$cumulative)) - (3*sd(as.numeric(traj_OA_summary$cumulative)))
Max_OAcum<-  mean(as.numeric(traj_OA_summary$cumulative)) + (3*sd(as.numeric(traj_OA_summary$cumulative)))
summary(traj_OA_summary$cumulative)

#test differences in cumulative movement in olfaction ablated toads
wilcox.test(cumulative~Ablation, traj_OA_summary)
wilcox.test(cumulative~Ablation, traj_OA_summary[traj_OA_summary$homing == 0,])
wilcox.test(cumulative~Ablation, traj_OA_summary[traj_OA_summary$homing == 1,])

#summarize cumulative movement in olfaction ablated toads
#homing
mean(traj_OA_summary[traj_OA_summary$homing == 1 & traj_OA_summary$Ablation == "OAC",]$cumulative)
sd(traj_OA_summary[traj_OA_summary$homing == 1 & traj_OA_summary$Ablation == "OAC",]$cumulative)
mean(traj_OA_summary[traj_OA_summary$homing == 1 & traj_OA_summary$Ablation == "OAE",]$cumulative)
sd(traj_OA_summary[traj_OA_summary$homing == 1 & traj_OA_summary$Ablation == "OAE",]$cumulative)
#non-homing
mean(traj_OA_summary[traj_OA_summary$homing == 0 & traj_OA_summary$Ablation == "OAC",]$cumulative)
sd(traj_OA_summary[traj_OA_summary$homing == 0 & traj_OA_summary$Ablation == "OAC",]$cumulative)
mean(traj_OA_summary[traj_OA_summary$homing == 0 & traj_OA_summary$Ablation == "OAE",]$cumulative)
sd(traj_OA_summary[traj_OA_summary$homing == 0 & traj_OA_summary$Ablation == "OAE",]$cumulative)

#no outliers, cumulative movement MA
Min_MAcum <- mean(as.numeric(traj_MA_summary$cumulative)) - (3*sd(as.numeric(traj_MA_summary$cumulative)))
Max_MAcum<-  mean(as.numeric(traj_MA_summary$cumulative)) + (3*sd(as.numeric(traj_MA_summary$cumulative)))
summary(traj_MA_summary$cumulative)

#test differences in cumulative movement in magnetoreception ablated toads
wilcox.test(cumulative~Ablation, traj_MA_summary)
wilcox.test(cumulative~Ablation, traj_MA_summary[traj_MA_summary$homing == 0,])
wilcox.test(cumulative~Ablation, traj_MA_summary[traj_MA_summary$homing == 1,])

#no straightness outliers
Min_OAstr <- mean(as.numeric(traj_OA_summary[traj_OA_summary$homing==1,]$straightness)) - (3*sd(as.numeric(traj_OA_summary[traj_OA_summary$homing==1,]$straightness)))
Max_OAstr<-  mean(as.numeric(traj_OA_summary[traj_OA_summary$homing==1,]$straightness)) + (3*sd(as.numeric(traj_OA_summary[traj_OA_summary$homing==1,]$straightness)))
Min_MAstr <- mean(as.numeric(traj_MA_summary[traj_MA_summary$homing==1,]$straightness)) - (3*sd(as.numeric(traj_MA_summary[traj_MA_summary$homing==1,]$straightness)))
Max_MAstr<-  mean(as.numeric(traj_MA_summary[traj_MA_summary$homing==1,]$straightness)) + (3*sd(as.numeric(traj_MA_summary[traj_MA_summary$homing==1,]$straightness)))

#test differences in straightness in ablated toads
wilcox.test(straightness~Ablation, traj_OA_summary)#ns
wilcox.test(straightness~Ablation, traj_OA_summary[traj_OA_summary$homing == 1,])#ns
wilcox.test(straightness~Ablation, traj_MA_summary)#ns
wilcox.test(straightness~Ablation, traj_MA_summary[traj_MA_summary$homing == 1,])#ns

#no duration outliers
Min_OAdur <- mean(as.numeric(traj_OA_summary[traj_OA_summary$homing==1,]$duration)) - (3*sd(as.numeric(traj_OA_summary[traj_OA_summary$homing==1,]$duration)))
Max_OAdur<-  mean(as.numeric(traj_OA_summary[traj_OA_summary$homing==1,]$duration)) + (3*sd(as.numeric(traj_OA_summary[traj_OA_summary$homing==1,]$duration)))
summary(traj_OA_summary[traj_OA_summary$homing==1,]$duration)
Min_MAdur <- mean(as.numeric(traj_MA_summary[traj_MA_summary$homing==1,]$duration)) - (3*sd(as.numeric(traj_MA_summary[traj_MA_summary$homing==1,]$duration)))
Max_MAdur<-  mean(as.numeric(traj_MA_summary[traj_MA_summary$homing==1,]$duration)) + (3*sd(as.numeric(traj_MA_summary[traj_MA_summary$homing==1,]$duration)))
summary(traj_MA_summary[traj_MA_summary$homing==1,]$duration)

#effect of ablation on homing duration
wilcox.test(duration~Ablation, traj_OA_summary[traj_OA_summary$homing == 1,])#ns
wilcox.test(duration~Ablation, traj_MA_summary[traj_MA_summary$homing == 1,])#ns

####################################################
#Circular Statistics and Plots

#Calculate Circular average fo absolute angles
traj_df$abs.angle_rad <- circular(traj_df$abs.angle, units = "radians")
#Calculate Circular average for rel angles
traj_df$rel.angle_rad <- circular(traj_df$rel.angle, units = "radians")
#make positive if neg
traj_df$abs.angle_rad[traj_df$abs.angle_rad < 0 & !is.na(traj_df$abs.angle_rad)] <- traj_df$abs.angle_rad[traj_df$abs.angle_rad < 0 & !is.na(traj_df$abs.angle_rad)] + 2 * pi
traj_df$rel.angle_rad[traj_df$rel.angle_rad < 0 & !is.na(traj_df$rel.angle_rad)] <- traj_df$rel.angle_rad[traj_df$rel.angle_rad < 0 & !is.na(traj_df$rel.angle_rad)] + 2 * pi

# Group by id and calculate circular mean for abs.angle_rad
for(i in sequence(length(traj_all_summary$id))){
  traj_all_summary$mean_abs_angle[i] <- mean.circular(!is.na(traj_df[traj_df$id == traj_all_summary$id[i],]$abs.angle_rad))}

# Group by id and calculate circular mean for rel.angle_rad
for(i in sequence(length(traj_all_summary$id))){
  traj_all_summary$mean_rel_angle[i] <- mean.circular(!is.na(traj_df[traj_df$id == traj_all_summary$id[i],]$rel.angle_rad))}

#for abs.angle, calculate the delta from the home angle
names(home_ang__df2 )[names(home_ang__df2 ) == 'abs.angle'] <- 'home.angle'
traj_all_summary <- left_join(traj_all_summary, home_ang__df2)
traj_all_summary$delta <- traj_all_summary$home.angle- traj_all_summary$mean_abs_angle

#make sircular summaries for homing and non-homing animals
circ.summary(traj_all_summary[traj_all_summary$homing ==1,]$delta)
circ.summary(traj_all_summary[traj_all_summary$homing ==0,]$delta)

circ.summary(traj_all_summary[traj_all_summary$homing ==1,]$mean_rel_angle)
circ.summary(traj_all_summary[traj_all_summary$homing ==0,]$mean_rel_angle)

#Looking at abs angle first
#subset for homing toads and format circular data
circ_home <- filter(traj_all_summary, homing == 1)
circ_home <- circular(circ_home$delta, units = "radians", zero = 0, rotation = "clock")

#test whether homing toads are oriented towards home
home<- circular(0, units = "radians")
rayleigh.test(circ_home, home) #significant towards home
rayleigh.test(circ_home)

#do the same with non-homing toads
circ_nh <- filter(traj_all_summary, homing == 0)
weird<- circular(pi/2, units = "radians")
circ_nh <- circular((circ_nh$delta), units = "radians", zero = 0, rotation = "clock")
rayleigh.test(circ_nh, home)
rayleigh.test(circ_nh, weird)
rayleigh.test(circ_nh)

# Manova approach for comparing two circular distributions based on Landler et al. 2021
manova_angles<-manova(cbind(cos(traj_all_summary$delta), sin(traj_all_summary$delta)) ~ traj_all_summary$homing)
summary (manova_angles)

#now look at relative angles
circ_home_rel <- filter(traj_all_summary, homing == 1)
circ_home_rel <- circular(circ_home_rel$mean_rel_angle, units = "radians", zero = 0, rotation = "clock")

home<- circular(0, units = "radians")
rayleigh.test(circ_home_rel, home)

circ_nh_rel <- filter(traj_all_summary, homing == 0)
circ_nh_rel <- circular((circ_nh_rel$mean_rel_angle), units = "radians", zero = 0, rotation = "clock")
rayleigh.test(circ_nh_rel, home)

manova_angles_rel<-manova(cbind(cos(traj_all_summary$mean_rel_angle), sin(traj_all_summary$mean_rel_angle)) ~ traj_all_summary$homing)
summary (manova_angles_rel)

## Circular plots

#plot data for homing toads
dev.off()

#set plot margins to zero and use "shrink" parameter to fit in all the points
par(mai=c(0.25, 0.25, 0.25, 0.25))
plot(circ_home, axes=FALSE, stack = TRUE, bin = 72, sep = 0.07, shrink = 1.5, main = "Homing")
ticks.circular(circular(seq(0,2*pi,pi/8)), tcl=0.075)
rose.diag(circ_home, bins=16, col="grey", cex=1.5, prop=1.3, add=TRUE)
arrows.circular(mean(circ_home), y = rho.circular(circ_home), lwd = 3)
lines(density.circular(circ_home, bw=30), lwd=2, lty=3)

#plot data for non-homing toads
#set plot margins to zero and use "shrink" parameter to fit in all the points
par(mai=c(0.25, 0.25, 0.25, 0.25))
plot(circ_nh, axes=FALSE, stack = TRUE, bin = 72, sep = 0.07, shrink = 1.5, main = "Nonhoming")
ticks.circular(circular(seq(0,2*pi,pi/8)), tcl=0.075)
rose.diag(circ_nh, bins=16, col="grey", cex=1.5, prop=1.3, add=TRUE)
arrows.circular(mean(circ_nh), y = rho.circular(circ_nh), lwd = 3)
lines(density.circular(circ_nh, bw=30), lwd=2, lty=3)

#rel angles homing
par(mai=c(0.25, 0.25, 0.25, 0.25))
plot(circ_home_rel, axes=FALSE, stack = TRUE, bin = 72, sep = 0.07, shrink = 1.5, main = "Homing")
ticks.circular(circular(seq(0,2*pi,pi/8)), tcl=0.075)
rose.diag(circ_home_rel, bins=16, col="grey", cex=1.5, prop=1.3, add=TRUE)
arrows.circular(mean(circ_home_rel), y = rho.circular(circ_home_rel), lwd = 3)
lines(density.circular(circ_home_rel, bw=30), lwd=2, lty=3)

#rel angles non-homing toads
#set plot margins to zero and use "shrink" parameter to fit in all the points
par(mai=c(0.25, 0.25, 0.25, 0.25))
plot(circ_nh_rel, axes=FALSE, stack = TRUE, bin = 72, sep = 0.07, shrink = 1.5, main = "Nonhoming")
ticks.circular(circular(seq(0,2*pi,pi/8)), tcl=0.075)
rose.diag(circ_nh_rel, bins=16, col="grey", cex=1.5, prop=1.3, add=TRUE)
arrows.circular(mean(circ_nh_rel), y = rho.circular(circ_nh_rel), lwd = 3)
lines(density.circular(circ_nh_rel, bw=30), lwd=2, lty=3)

###############################################################################
#incorporating weather data
weather <- read.csv("~/Documents/DocumentsDanielWork/HI_2022/Weather.csv")

kruskal.test(PRCP~loc,weather)
dunnTest(PRCP~loc,weather, method = "bh")
boxplot(PRCP~loc,weather)
kruskal.test(TMAX~loc,weather)
dunnTest(TMAX~loc,weather, method = "bh")
boxplot(TMAX~loc,weather)
kruskal.test(TMIN~loc,weather)
dunnTest(TMIN~loc,weather, method = "bh")
boxplot(TMIN~loc,weather)
####################################################################################################
#PART 2
#Analysis of Baseline Tracking Data
####################################################################################################
#read the spreadsheets into r
#baseline tracking coordinates
Baseline <- read.csv("~/Documents/DocumentsDanielWork/HI_2022/ArcGIS Exports/Baseline.csv")
#infoabout all toads
toad_info_2 <- read.csv("~/Documents/DocumentsDanielWork/HI_2022/ArcGIS Exports/HI 2022 Toad List.csv")

#convert time-stamps to POSIXct objects for the trajectory
da_b <- as.POSIXct(strptime(Baseline$Timestamp_E,"%Y-%m-%d %H:%M"))

#Fix because two values being returned as NA for some reason
na_indices <- which(is.na(da_b))
print(na_indices)
da_b[451] <- as.POSIXct("2022-03-27 02:53", format = "%Y-%m-%d %H:%M", tz = "UTC")
da_b[549] <- as.POSIXct("2022-03-27 02:58", format = "%Y-%m-%d %H:%M", tz = "UTC")

#make trajectory items
base_traj <- as.ltraj(xy=Baseline[,c("POINT_X","POINT_Y")], date = da_b, id = Baseline$ID)

#turn into dataframes
base_df <- ld(base_traj)
base_df$index <- as.numeric(row.names(base_df))
order(base_df$index)

#make summaries and calculate duration
base_summary <-summary(base_traj)
base_summary <- right_join(base_summary, toad_info_2)

#baseline tracking duration in days
base_summary$base.traj.time <- base_summary$date.end - base_summary$date.begin
#baseline tracking duration in hours
base_summary$base.traj.hrs <- as.numeric(base_summary$base.traj.time*24)
base_df <- left_join(base_df, base_summary[,c("id","Location")])

#total distance moved over observed time 
for (i in sequence(length(base_summary$id))){
  traj_sub <- base_traj[which(id(base_traj) == base_summary$id[i])]
  base_summary$base.cumulative[i] = sum(na.omit(traj_sub[[1]]$dist))
}

# net displacement of observed period, and straightness
for (i in sequence(length(base_summary$id))){
  traj_sub <- base_traj[which(id(base_traj) == base_summary$id[i])]
  x <- traj_sub[[1]]$x
  y <- traj_sub[[1]]$y
  lngt = length(traj_sub[[1]]$x)
  base_summary$base.return_transect[i] = pointDistance(c(x[1],y[1]),c(x[lngt],y[lngt]),lonlat = FALSE, allpairs =FALSE)
}

base_summary$base.straightness <- (base_summary$base.return_transect/base_summary$base.cumulative)

#movement range of baseline 
for (i in sequence(length(base_summary$id)))
{
  sub <- base_df[base_df$id == base_summary$id[i],]
  dists <- c()
  for(j in sequence(length(sub$x)-1)){
    for (z in seq(j+1,  length(sub$x))){
      d <- as.numeric(pointDistance(c(sub$x[j],sub$y[j]),c(sub$x[z],sub$y[z]),lonlat = FALSE, allpairs =FALSE))
      dists <- append(dists, d)
    }
  }
  base_summary$base.range[i] <- max(dists)
}

#average distance moved per day
base_summary$avg_dist <- as.numeric(base_summary$base.cumulative)/as.numeric(base_summary$base.traj.time)
###########################################################################################################

#organize dataframe and select only toads tracked more than 3 days
base_summary <- base_summary[!is.na(as.numeric(base_summary$base.traj.time)) & as.numeric(base_summary$base.traj.time) > 2.5, ]

base_summary <- as_tibble(base_summary) %>%
  dplyr::select(id = id, sex = Sex, mass = Mass, svl = SVL, loc = Location, base.traj.time = base.traj.time, base.traj.hrs = base.traj.hrs, base.cumulative=base.cumulative, base.return_transect=base.return_transect, base.straightness=base.straightness, base.range=base.range, avg_dist=avg_dist) %>%
  arrange(id) %>%
  group_by(id)

#for reporting  information in paper
mean(base_summary$mass)
sd(base_summary$mass)
mean(base_summary$svl)
sd(base_summary$svl)

#mass not normal
shapiro.test(base_summary$mass)
#svl normal
shapiro.test(base_summary$svl)

#compare mass across tracking sites
kruskal.test(mass~loc, base_summary)
dunnTest(mass ~ loc, data=base_summary)

#compare svl across tracking sites
a <- aov(svl~loc, base_summary)
summary(a)
a_posthoc <- TukeyHSD(a)
a_posthoc

#compare svl and mass by sec
t.test(svl~sex, base_summary)
wilcox.test(mass~sex, data = base_summary, exact = FALSE)

##########################################################################

#remove three toads whose movements not fully accounted for
base_summary <- base_summary[as.numeric(base_summary$base.traj.time)<20,] 

#movement range not normal
shapiro.test(base_summary$base.range)

#Check range for outliers- there are three
Min_range <- mean(as.numeric(base_summary$base.range)) - (3*sd(as.numeric(base_summary$base.range)))
Max_range<-  mean(as.numeric(base_summary$base.range)) + (3*sd(as.numeric(base_summary$base.range)))

#compare range across sites when removing outliers
kruskal.test(base.range~loc, base_summary[base_summary$base.range < Max_range,])

#compare range across sites without removing outliers
kruskal.test(base.range~loc, base_summary)
dunnTest(base.range ~ loc, data=base_summary)

#compare range across sexes when removing outliers
wilcox.test(base.range~sex, base_summary[base_summary$base.range < Max_range,])
#compare range across sexes without removing outliers
wilcox.test(base.range~sex, base_summary)

#summaries for reporting in paper
summary(base_summary$base.range)
sd(base_summary$base.range)

summary(base_summary$base.cumulative)
sd(base_summary$base.cumulative)

summary(base_summary$avg_dist)
sd(base_summary$avg_dist)

summary(as.numeric(base_summary$base.traj.time))
sd(as.numeric(base_summary$base.traj.time))

#model range as a function of size
a <- lm(base.range~svl, base_summary)
summary(a)

a2 <- lmer(base.range ~ svl + (1|loc), base_summary)
summary(a2)
#############################################################################################################
#PART 3
#Script for looking at effects of ablations on pre-translocation movements
#############################################################################################################
#read the spreadsheets into r
#Baseline coordinates for ablation toads
Ab_base <- read.csv("~/Documents/DocumentsDanielWork/HI_2022/ArcGIS Exports/Ablation_Baseline.csv")
#Info for translocated toads
toad_info <- read.csv("~/Documents/DocumentsDanielWork/HI_2022/ArcGIS Exports/TransToadInfo.csv")
#Info for all toads
toad_info_2 <- read.csv("~/Documents/DocumentsDanielWork/HI_2022/ArcGIS Exports/HI 2022 Toad List.csv")

#Split up baseline coordinates into pre- and post-ablation 
before <- Ab_base[Ab_base$Pre.post == 0,]
after <-  Ab_base[Ab_base$Pre.post == 1,]

#convert the timestamps to a POSIXct object so it cab be used in adehabitatlt
da_before <- as.POSIXct(strptime(before$Timestamp_E,"%Y-%m-%d %H:%M"))
da_after <- as.POSIXct(strptime(after$Timestamp_E,"%Y-%m-%d %H:%M"))

#Fix because two values being returned as NA for some reason
na_indices <- which(is.na(da_after))
print(na_indices)
da_after[43] <- as.POSIXct("2022-03-27 02:53", format = "%Y-%m-%d %H:%M", tz = "UTC")
da_after[81] <- as.POSIXct("2022-03-27 02:58", format = "%Y-%m-%d %H:%M", tz = "UTC")


#make trajectory items 
before_traj <- as.ltraj(xy=before[,c("POINT_X","POINT_Y")], date = da_before, id = before$ID)
after_traj <- as.ltraj(xy=after[,c("POINT_X","POINT_Y")], date = da_after, id = after$ID)

#turn into dataframes
before_df <- ld(before_traj)
after_df <- ld(after_traj)

#reestablish pre/post-ablation coding
before_df$prepost <- 0
after_df$prepost <- 1

#make summaries
before_summary <-summary(before_traj)
before_summary$prepost <- 0
after_summary <-summary(after_traj)
after_summary$prepost <- 1

#calculate cumulative distance traveled
for (i in sequence(length(before_summary$id))){
  traj_sub <- before_traj[which(id(before_traj) == before_summary$id[i])]
  before_summary$base.cumulative2[i] = sum(na.omit(traj_sub[[1]]$dist))
}

for (i in sequence(length(after_summary$id))){
  traj_sub <- after_traj[which(id(after_traj) == after_summary$id[i])]
  after_summary$base.cumulative2[i] = sum(na.omit(traj_sub[[1]]$dist))
}

#Calculate displacement from beginning to end of each period, and straightness
for (i in sequence(length(before_summary$id))){
  traj_sub <- before_traj[which(id(before_traj) == before_summary$id[i])]
  x <- traj_sub[[1]]$x
  y <- traj_sub[[1]]$y
  lngt = length(traj_sub[[1]]$x)
  before_summary$base.return_transect[i] = pointDistance(c(x[1],y[1]),c(x[lngt],y[lngt]),lonlat = FALSE, allpairs =FALSE)
}

for (i in sequence(length(after_summary$id))){
  traj_sub <- after_traj[which(id(after_traj) == after_summary$id[i])]
  x <- traj_sub[[1]]$x
  y <- traj_sub[[1]]$y
  lngt = length(traj_sub[[1]]$x)
  after_summary$base.return_transect[i] = pointDistance(c(x[1],y[1]),c(x[lngt],y[lngt]),lonlat = FALSE, allpairs =FALSE)
}

before_summary$base.straightness <- (before_summary$base.return_transect/before_summary$base.cumulative)
after_summary$base.straightness <- (after_summary$base.return_transect/after_summary$base.cumulative)

#calculate movement ranges for each period
for (i in sequence(length(before_summary$id)))
{
  sub <- before_df[before_df$id == before_summary$id[i],]
  dists <- c()
  for(j in sequence(length(sub$x)-1)){
    for (z in seq(j+1,  length(sub$x))){
      d <- as.numeric(pointDistance(c(sub$x[j],sub$y[j]),c(sub$x[z],sub$y[z]),lonlat = FALSE, allpairs =FALSE))
      dists <- append(dists, d)
    }
  }
  before_summary$base.range[i] <- max(dists)
}

for (i in sequence(length(after_summary$id)))
{
  sub <- after_df[after_df$id == after_summary$id[i],]
  dists <- c()
  for(j in sequence(length(sub$x)-1)){
    for (z in seq(j+1,  length(sub$x))){
      d <- as.numeric(pointDistance(c(sub$x[j],sub$y[j]),c(sub$x[z],sub$y[z]),lonlat = FALSE, allpairs =FALSE))
      dists <- append(dists, d)
    }
  }
  after_summary$base.range[i] <- max(dists)
}

#consolidate dataframes
ab_base_df <- rbind(after_df, before_df)
ab_base_summary <- rbind(before_summary,after_summary)

ab_base_summary <- left_join(ab_base_summary, toad_info_2)
ab_base_summary <- left_join(ab_base_summary, toad_info[,c("id", "Pair")])

#calculate time of each period and proportion of points post ablation in minimum
#convex polygon encompassing pe-ablation points (calculated in ArcGIS)
ab_base_summary$base.traj.time <- ab_base_summary$date.end - ab_base_summary$date.begin
ab_base_summary$base.traj.hrs <- as.numeric(ab_base_summary$base.traj.time*24)
ab_base_summary$post_proportion <- ab_base_summary$PostPtsinPreMCP/ab_base_summary$PostPts

#split up into olfactory and magnetoreception
olf_base_summary <- ab_base_summary[ab_base_summary$Ablation == "OAC" | ab_base_summary$Ablation == "OAE", ]
ol_before <- olf_base_summary[olf_base_summary$prepost == 0,]
ol_after <- olf_base_summary[olf_base_summary$prepost == 1,]
mag_base_summary <- ab_base_summary[ab_base_summary$Ablation == "MAC" | ab_base_summary$Ablation == "MAE", ]
mag_before <- mag_base_summary[mag_base_summary$prepost == 0,]
mag_after <- mag_base_summary[mag_base_summary$prepost == 1,]
shapiro.test(ab_base_summary$base.range)

#2 outliers
Min_OArange <- mean(olf_base_summary$base.range) - (3*sd(olf_base_summary$base.range))
Max_OArange<-  mean(olf_base_summary$base.range) + (3*sd(olf_base_summary$base.range))
summary(olf_base_summary$base.range)

# 2 outliers
Min_MArange <- mean(mag_base_summary$base.range) - (3*sd(mag_base_summary$base.range))
Max_MArange<-  mean(mag_base_summary$base.range) + (3*sd(mag_base_summary$base.range))
summary(mag_base_summary$base.range)

#COMPARIONS REPORTED IN PAPER

#olfaction
#ns-  range in control animals
wilcox.test(as.numeric(ol_before[ol_before$Ablation=="OAC",]$base.range), as.numeric(ol_after[ol_after$Ablation=="OAC",]$base.range), paired = TRUE, conf.int = T, conf.level = .95)

#ns - range in experimental animals
wilcox.test(as.numeric(ol_before[ol_before$Ablation=="OAE",]$base.range), as.numeric(ol_after[ol_after$Ablation=="OAE",]$base.range), paired = TRUE, conf.int = T, conf.level = .95)

#similarity before and after-  significantly different
wilcox.test(post_proportion~Ablation, olf_base_summary, exact = FALSE)

#magnetoreception
#sig p = 0.04
wilcox.test(as.numeric(mag_before[mag_before$Ablation=="MAC",]$base.range), as.numeric(mag_after[mag_after$Ablation=="MAC",]$base.range), paired = TRUE, conf.int = T, conf.level = .95)

#sig, p = 0.007
wilcox.test(as.numeric(mag_before[mag_before$Ablation=="MAE",]$base.range), as.numeric(mag_after[mag_after$Ablation=="MAE",]$base.range), paired = TRUE, conf.int = T, conf.level = .95)

#remove outliers- significance holds
mag_before2<- mag_before[mag_before$id != "B8",]
mag_before2<- mag_before2[mag_before2$id != "D6",]
mag_after2<- mag_after[mag_after$id != "B8",]
mag_after2<- mag_after2[mag_after2$id != "D6",]
wilcox.test(as.numeric(mag_before2[mag_before2$Ablation=="MAE",]$base.range), as.numeric(mag_after2[mag_after2$Ablation=="MAE",]$base.range), paired = TRUE, conf.int = T, conf.level = .95)

#similarity before and after-  ns
wilcox.test(post_proportion~Ablation, mag_base_summary, exact = FALSE)
#######################################################################################################
#PART 4
#Analysis of pS6 count data
######################################################################################################

#THIS SECTION READS IN ALL THE SPREADSHEETS AND MAKES ALTERATIONS TO THE DATAFRAMES
################################################################################
#read in spreadhseets of counting brain toad info
counts <- read.csv("~/Documents/DocumentsDanielWork/HI_2022/Brain Info/BrainCounts_02.csv")
brains <- read.csv("~/Documents/DocumentsDanielWork/HI_2022/Brain Info/Brains.csv")
toads <-  read.csv("~/Documents/DocumentsDanielWork/HI_2022/Brain Info/Toads.csv")
olf_ct <-  read.csv("~/Documents/DocumentsDanielWork/HI_2022/Brain Info/olf_ct.csv")
#generated in HI_Toads_spatial_Data_053024.R
traj_all_summary <- read.csv("~/Documents/DocumentsDanielWork/HI_2022/traj_all_summary.csv")

#brains with no counts
brains <- brains[brains$Toad != "C3",]
brains <- brains[brains$Toad != "C8",]
brains <- brains[brains$Toad != "D10",]
brains <- brains[brains$Toad != "F9",]

lost <- brains[brains$Condition == "Lost",]
#remove *Homing from counts and brains (brains collected at incorrect timepoint)
counts <- counts[counts$Toad != "G3",]
counts <- counts[counts$Toad != "F7",]
brains <- brains[brains$Toad != "G3",]
brains <- brains[brains$Toad != "F7",]

#clean up
toads <- toads[,c("Toad", "Sex", "Mass", "SVL", "Location")] 
counts <- counts[,c("Toad", "Slice", "Brain.Region", "Count", "Area")]

#make septum measurement
counts_new <- counts %>%
  # Filter for Ms and Ls
  filter(Brain.Region %in% c("Ms", "Ls")) %>%
  # Group by Toad and Slice
  group_by(Toad, Slice) %>%
  # Summarize Count and Area for Ms and Ls
  summarise(
    Brain.Region = "SepNew", 
    Count = sum(Count),
    Area = sum(Area),
    .groups = 'drop'  # Ungroup after summarizing
  )

counts <- rbind(counts,counts_new)
counts <- counts[counts$Brain.Region != "Sep",]
counts <- counts[counts$Brain.Region != "SepNew",]

#Need to remove slices 1-3 for Dp, Sep, Stri, as these only had counts in 6 sections
list <- c()
for (i in sequence(length(counts$Toad))){
  if ((counts$Brain.Region[i]=="DP" | counts$Brain.Region[i]=="Sep" | counts$Brain.Region[i]=="Stri") & (counts$Slice[i] < 4)){
    list <- append(list,i)
  }
}
counts <- counts[-list,]

#fill in non counted cells with 0s
counts[is.na(counts)] <- 0

counts<- right_join(brains, counts, by="Toad")
counts <- right_join (toads, counts, by = "Toad")

#convert Area from micrometers sq to mm sq and calculate density
counts$Areamm <- as.numeric(counts$Area)/1000000
counts$density <- as.numeric(counts$Count)/as.numeric(counts$Areamm)

#at least temporarary- remove cMP/vMP from counts- not considered in analysis
counts <- counts[counts$Brain.Region != "cMP",]
counts <- counts[counts$Brain.Region != "vMP",]

for (i in sequence(length(counts$Toad))){
  if (counts$Slice[i] < 4){counts$sections[i] <- "Anterior"}
  if (counts$Slice[i] > 3 && counts$Slice[i] < 7 ){counts$sections[i] <- "Medial"}
  if (counts$Slice[i] > 6){counts$sections[i] <- "Posterior"}
}

####THIS SECTION DEALS WITH OLF ABLATION OLF BULBS- comparing pS6 count in olfactory bukb
#between control and experimental 
################################################################################

olf_ct <- left_join(olf_ct, traj_all_summary[,c("id","Ablation")])

olf_ct_means <- olf_ct %>%
  group_by(id)%>%
  summarize(pS6 = mean(count), density = mean(density), Ablation = first(Ablation))%>%
  droplevels()
olf_ct_means


wilcox.test(density~Ablation, olf_ct, excact= FALSE)
wilcox.test(pS6~Ablation, olf_ct_means, excact= FALSE)

#THIS SECTION CHECKS THE DISTRIBUTION OF DATA/ DOES SOME DATA EXPLORATION
################################################################################
#ccheck model fit
ggplot(counts, aes(y=density, x=Brain.Region, fill=Condition)) + geom_boxplot()

counts.lm = lm(density ~ Condition, data=counts) 
counts.stdres = rstandard(counts.lm)
summary(counts.lm)
AIC(counts.lm)

qqp(counts.stdres, "norm")
qqp(counts.stdres, "lnorm")

nb <- counts$Count + 1

#THIS SEEMS TO LOOK GOOD negative binomial fit
nbinom <- fitdistr(nb, "Negative Binomial")
qqp(nb, "nbinom", size = nbinom$estimate[[1]], mu = nbinom$estimate[[2]])

poisson <- fitdistr(nb, "Poisson")
qqp(nb, "pois", lambda = poisson$estimate)

gamma <- fitdistr(nb, "gamma")
qqp(nb, "gamma", shape = gamma$estimate[[1]], rate = gamma$estimate[[2]])

#THIS SECTION IS FOR MAKING MODELS OF THE COUNT DATA
################################################################################

#model all toads, with homing condition and brain region and their interaction as fixed effects, Toad ID as random effect, and brain area as offset
m1 <-  glmmTMB(Count ~ Condition*Brain.Region + (1|Toad) + offset(log(as.integer(Area))), family=nbinom2(link="log"), data=counts)
summary(m1) 
glmmTMB:::Anova.glmmTMB(m1, type="II")
AIC(m1)
#pairwise comparisons
emmeans(m1, specs = pairwise ~ Condition|Brain.Region, adjust="tukey")

#next three models splitting brain into anterior, medial, posterior, to look at differences in the Mp across the AP axi
anterior <- glmmTMB(Count ~ Condition*Brain.Region + (1|Toad) + offset(log(as.integer(Area))), family=nbinom2(link="log"), data=counts[counts$sections == "Anterior",])
summary(anterior)
glmmTMB:::Anova.glmmTMB(anterior, type="II")
AIC(anterior)
#pairwise comparisons
emmeans(anterior, specs = pairwise ~ Condition|Brain.Region, adjust="tukey")

medial <- glmmTMB(Count ~ Condition*Brain.Region + (1|Toad) + offset(log(as.integer(Area))), family=nbinom2(link="log"), data=counts[counts$sections == "Medial",])
summary(medial)
glmmTMB:::Anova.glmmTMB(medial, type="II")
AIC(medial)
#pairwise comparisons
emmeans(medial, specs = pairwise ~ Condition|Brain.Region, adjust="tukey")

posterior <- glmmTMB(Count ~ Condition*Brain.Region + (1|Toad) + offset(log(as.integer(Area))), family=nbinom2(link="log"), data=counts[counts$sections == "Posterior",])
summary(posterior)
glmmTMB:::Anova.glmmTMB(posterior, type="II")
AIC(posterior)
#pairwise comparisons
emmeans(posterior, specs = pairwise ~ Condition|Brain.Region, adjust="tukey")

#model that included sex as a predictor (no significant effect)
m2 <-  glmmTMB(Count ~ Condition*Brain.Region*Sex + (1|Toad) + offset(log(as.integer(Area))), family=nbinom2(link="log"), data=counts)
summary(m2) 
glmmTMB:::Anova.glmmTMB(m2, type="II")
AIC(m2)
#pairwise comparisons
emmeans(m2, specs = pairwise ~ Sex|Brain.Region*Condition, adjust="tukey")
emmeans(m2, specs = pairwise ~ Condition|Brain.Region, adjust="tukey")

#model that includes ablation as a predictor
#modifications- take out baseline animals and MAE* animals
counts2 <- counts[counts$Ablation != "MAE*",]
counts2 <-  counts2[counts2$Condition != "Baseline",]
m3 <-  glmmTMB(Count ~ Condition*Brain.Region*Ablation + (1|Toad) + offset(log(as.integer(Area))), family=nbinom2(link="log"), data=counts2)
summary(m3) 
glmmTMB:::Anova.glmmTMB(m3, type="II")
AIC(m3)
#pairwise comparisons
emmeans(m3, specs = pairwise ~ Ablation|Brain.Region, adjust="tukey")
emmeans(m3, specs = pairwise ~ Ablation|Brain.Region*Condition, adjust="tukey")
emmeans(m3, specs = pairwise ~ Condition|Brain.Region, adjust="tukey")

#modeling only nonablated toads with homing condition and brain region and their interaction as fixed effects, Toad ID as randome effect, and brain area as offset
unablated = counts[counts$Ablation == "None",]
m1x <-  glmmTMB(Count ~ Condition*Brain.Region + (1|Toad) + offset(log(as.integer(Area))), family=nbinom2(link="log"), data=unablated)
summary(m1x) 
glmmTMB:::Anova.glmmTMB(m1x, type="II")
AIC(m1x)
#pairwise comparisons
emmeans(m1x, specs = pairwise ~ Condition|Brain.Region, adjust="tukey")

#modeling only non-homing
lost <- counts[counts$Condition == "Lost",]

lost_model <- glmmTMB(Count ~ Location*Brain.Region + (1|Toad) + offset(log(as.integer(Area))), family=nbinom2(link="log"), data=lost)
summary(lost_model)
glmmTMB:::Anova.glmmTMB(lost_model, type="II")
emmeans(lost_model, specs = pairwise ~ Location|Brain.Region, adjust="tukey")

#make a simplified version of the dataframe with mean count for each brain region for each toad
ps6_means <- counts %>%
  group_by(Toad, Brain.Region)%>%
  mutate(density = Count/Areamm)%>%
  summarize(group = first(Condition), pS6 = mean(Count), density = mean(density), Ablation = first(Ablation))%>%
  droplevels()
ps6_means

#make a simplified version of the dataframe with mean count for each brain region for each toad for the model without baseline toads
ps6_means2 <- counts2 %>%
  group_by(Toad, Brain.Region)%>%
  mutate(density = Count/Areamm)%>%
  summarize(group = first(Ablation), pS6 = mean(Count), density = mean(density), Condition = first(Condition))%>%
  droplevels()
ps6_means2
##############################################################################
#COMPARING PERFORMANCE BY STRAIGHTNESS, DURATION

#look at only translocated toads
translocated <- ps6_means %>%
  filter(group != "Baseline")

#add straightness values and duration values to count information
strght <- data.frame(Toad = traj_all_summary$id, straightness = traj_all_summary$straightness, duration = traj_all_summary$duration, homing = traj_all_summary$homing, loc = traj_all_summary$loc )
translocated <- left_join(translocated, strght)
counts <- left_join(counts, strght)

#split into homing and not homing
translocated_homing <- translocated[translocated$homing == 1,]
translocated_lost <- translocated[translocated$homing == 0,]

#also look at count data for homing animals
homing_counts <- counts[counts$homing == 1,]

#Making a mixed linear effects model to show the correlation of brain activity by region and path straightnress
model_straightness <- lmer(Count ~ straightness * Brain.Region + (1|Toad), data = homing_counts)
summary(model_straightness)

#Now modeling activity individually for each brain region
#ns
stri_straightness <- lmer(Count ~ straightness + (1|Toad) + offset(log(as.integer(Area))), data = homing_counts[homing_counts$Brain.Region == "Stri",])
summary(stri_straightness)

#ns
mp_straightness <- lmer(Count ~ straightness + (1|Toad)+ offset(log(as.integer(Area))), data = homing_counts[homing_counts$Brain.Region == "MP",])
summary(mp_straightness)

#ns
dp_straightness <- lmer(Count ~ straightness + (1|Toad)+ offset(log(as.integer(Area))), data = homing_counts[homing_counts$Brain.Region == "DP",])
summary(dp_straightness)

#ns
sep_straightness <- lmer(Count ~ straightness + (1|Toad)+ offset(log(as.integer(Area))), data = homing_counts[homing_counts$Brain.Region == "SepNew",])
summary(sep_straightness)

msep_straightness <- lmer(Count ~ straightness + (1|Toad)+ offset(log(as.integer(Area))), data = homing_counts[homing_counts$Brain.Region == "Ms",])
summary(msep_straightness)

lsep_straightness <- lmer(Count ~ straightness + (1|Toad)+ offset(log(as.integer(Area))), data = homing_counts[homing_counts$Brain.Region == "Ls",])
summary(lsep_straightness)

#ns
lp_straightness <- lmer(Count ~ straightness + (1|Toad)+ offset(log(as.integer(Area))), data = homing_counts[homing_counts$Brain.Region == "LP",])
summary(lp_straightness)

#Making a mixed linear effects model to show the correlation of brain activity by region and homing duration

model_duration <- lmer(Count ~ duration * Brain.Region + (1|Toad) + offset(log(as.integer(Area))), data = homing_counts)
summary(model_duration)

#Now modeling activity individually for each brain region
#ns
stri_duration <- lmer(Count ~ duration + (1|Toad)+ offset(log(as.integer(Area))), data = homing_counts[homing_counts$Brain.Region == "Stri",])
summary(stri_duration)

#ns
mp_duration <- lmer(Count ~ duration + (1|Toad)+ offset(log(as.integer(Area))), data = homing_counts[homing_counts$Brain.Region == "MP",])
summary(mp_duration)

#ns
dp_duration <- lmer(Count ~ duration + (1|Toad)+ offset(log(as.integer(Area))), data = homing_counts[homing_counts$Brain.Region == "DP",])
summary(dp_duration)

#sig
sep_duration <- lmer(Count ~ duration + (1|Toad)+ offset(log(as.integer(Area))), data = homing_counts[homing_counts$Brain.Region == "SepNew",])
summary(sep_duration)

msep_duration <- lmer(Count ~ duration + (1|Toad)+ offset(log(as.integer(Area))), data = homing_counts[homing_counts$Brain.Region == "Ms",])
summary(msep_duration)

lsep_duration <- lmer(Count ~ duration + (1|Toad)+ offset(log(as.integer(Area))), data = homing_counts[homing_counts$Brain.Region == "Ls",])
summary(lsep_duration)

#ns
lp_duration <- lmer(Count ~ duration + (1|Toad)+ offset(log(as.integer(Area))), data = homing_counts[homing_counts$Brain.Region == "LP",])
summary(lp_duration)
###########################################################################################################
#PART 5
##Normalize trajectories and plot with origin being translocation release site
########################################################################################################
#remove unnecessary columns
toads_df$X = NULL

#normalize x and y coordinates with respect to translocation release point
toads_df <- toads_df %>%
  group_by(id) %>%
  mutate(x_norm = x_utm - x_utm[1]) %>%
  mutate(y_norm = y_utm - y_utm[1])

#add the home angles to the dataframe
toads_df <- right_join(toads_df, home_ang__df2)

#make the angles positive if negative
toads_df %>%
  mutate(angle.norm = ifelse(home.angle > 0, home.angle, 2*pi - abs(home.angle))) -> toads_df

#rotate with respect to the home angle to home is the positive y direction
toads_df$x_new <- toads_df$x_norm * cos((0) + (toads_df$angle.norm)) - toads_df$y_norm * sin((0) + (toads_df$angle.norm))
toads_df$y_new <- toads_df$y_norm * cos((0) + (toads_df$angle.norm)) + toads_df$x_norm * sin((0) + (toads_df$angle.norm))

#add ablation and distance to these dataframes
toads_df <- left_join(toads_df, brains2)

#plot  in circles
#Figure 2A
#all that were translocated 500
ggplot() + 
  geom_circle(aes(x0 = 0, y0 = 500, r = 200), linetype = "solid", color ="grey", fill = "grey", size = 0.25, alpha = 0.5) +
  geom_circle(aes(x0 = 0, y0 = 0, r = 100), linetype = "solid", size = 0.25, color = "grey") +
  geom_circle(aes(x0 = 0, y0 = 0, r = 200), linetype = "solid", size = 0.25, color = "grey") +
  geom_circle(aes(x0 = 0, y0 = 0, r = 300), linetype = "solid", size = 0.25, color = "grey") +
  geom_circle(aes(x0 = 0, y0 = 0, r = 400), linetype = "solid", size = 0.25, color = "grey") +
  geom_circle(aes(x0 = 0, y0 = 0, r = 500), linetype = "solid", size = 0.25, color = "grey") +
  geom_circle(aes(x0 = 0, y0 = 0, r = 600), linetype = "solid", size = 1, color = "black") +
  geom_segment(aes(x = -600, xend = 600, y = 0, yend = 0), linetype = "solid", size = 0.25, color = "grey") +
  geom_segment(aes(x = 0, xend = 0, y = -600, yend = 600), linetype = "solid", size = 0.25, color = "grey") + 
  geom_path(data = toads_df[(toads_df$Distance ==500 & toads_df$homing == 1),], aes(x = x_new, y = y_new, group = id, color = as.factor(homing)), arrow = arrow(angle = 20, length = unit(0.1, "inches"), ends = "last", type ="closed")) +
  geom_path(data = toads_df[(toads_df$Distance ==500 & toads_df$homing == 0),], aes(x = x_new, y = y_new, group = id, color = as.factor(homing)), arrow = arrow(angle = 20, length = unit(0.1, "inches"), ends = "last", type ="closed")) +
  coord_fixed(ratio = 1)+
  scale_color_manual(values = c("blue", "goldenrod")) +
  theme(
    panel.border = element_blank(),
    axis.text = element_blank(),
    axis.title.x=element_blank(),
    axis.title.y=element_blank(),
    panel.grid = element_blank(),
    axis.ticks = element_blank(),
    plot.background = element_rect(fill = "transparent"),
    panel.background = element_rect(fill = "transparent"),
    legend.position = "right",
    legend.title=element_blank())

#Figure 2B
#all that were translocated 1000 
ggplot() + 
  geom_circle(aes(x0 = 0, y0 = 1000, r = 200), linetype = "solid", color ="grey", fill = "grey", size = 0.25, alpha = 0.5) +
  geom_circle(aes(x0 = 0, y0 = 0, r = 200), linetype = "solid", size = 0.25, color = "grey") +
  geom_circle(aes(x0 = 0, y0 = 0, r = 400), linetype = "solid", size = 0.25, color = "grey") +
  geom_circle(aes(x0 = 0, y0 = 0, r = 600), linetype = "solid", size = 0.25, color = "grey") +
  geom_circle(aes(x0 = 0, y0 = 0, r = 800), linetype = "solid", size = 0.25, color = "grey") +
  geom_circle(aes(x0 = 0, y0 = 0, r = 1000), linetype = "solid", size = 0.25, color = "grey") +
  geom_circle(aes(x0 = 0, y0 = 0, r = 1200), linetype = "solid", size = 1, color = "black") +
  geom_segment(aes(x = -1200, xend = 1200, y = 0, yend = 0), linetype = "solid", size = 0.25, color = "grey") +
  geom_segment(aes(x = 0, xend = 0, y = -1200, yend = 1200), linetype = "solid", size = 0.25, color = "grey") + 
  geom_path(data = toads_df[toads_df$Distance ==1000,], aes(x = x_new, y = y_new, group = id, color = as.factor(homing)), arrow = arrow(angle = 20, length = unit(0.1, "inches"), ends = "last", type ="closed")) +
  coord_fixed(ratio = 1)+
  scale_color_manual(values = c("goldenrod", "blue")) +
  theme(
    panel.border = element_blank(),
    axis.text = element_blank(),
    axis.title.x=element_blank(),
    axis.title.y=element_blank(),
    panel.grid = element_blank(),
    axis.ticks = element_blank(),
    plot.background = element_rect(fill = "transparent"),
    panel.background = element_rect(fill = "transparent"),
    legend.position = ("right"),
    legend.title=element_blank())

#####Figure 3A - OLFACTORY
#olfactory that were translocated 1000
OA_df <- toads_df[toads_df$Ablation == "OAE" | toads_df$Ablation == "OAC",]
ggplot() + 
  geom_circle(aes(x0 = 0, y0 = 1000, r = 200), linetype = "solid", color ="grey", fill = "grey", size = 0.25, alpha = 0.5) +
  geom_circle(aes(x0 = 0, y0 = 0, r = 200), linetype = "solid", size = 0.25, color = "grey") +
  geom_circle(aes(x0 = 0, y0 = 0, r = 400), linetype = "solid", size = 0.25, color = "grey") +
  geom_circle(aes(x0 = 0, y0 = 0, r = 600), linetype = "solid", size = 0.25, color = "grey") +
  geom_circle(aes(x0 = 0, y0 = 0, r = 800), linetype = "solid", size = 0.25, color = "grey") +
  geom_circle(aes(x0 = 0, y0 = 0, r = 1000), linetype = "solid", size = 0.25, color = "grey") +
  geom_circle(aes(x0 = 0, y0 = 0, r = 1200), linetype = "solid", size = 1, color = "black") +
  geom_segment(aes(x = -1200, xend = 1200, y = 0, yend = 0), linetype = "solid", size = 0.25, color = "grey") +
  geom_segment(aes(x = 0, xend = 0, y = -1200, yend = 1200), linetype = "solid", size = 0.25, color = "grey") + 
  geom_path(data = OA_df[(OA_df$Distance ==1000 & OA_df$homing == 1),], aes(x = x_new, y = y_new, group = id, color = as.factor(Ablation)), arrow = arrow(angle = 20, length = unit(0.1, "inches"), ends = "last", type ="closed")) +
  geom_path(data = OA_df[(OA_df$Distance ==1000 & OA_df$homing == 0),], aes(x = x_new, y = y_new, group = id, color = as.factor(Ablation)), arrow = arrow(angle = 20, length = unit(0.1, "inches"), ends = "last", type ="closed")) +
  coord_fixed(ratio = 1)+
  scale_color_manual(values = c("darkgreen", "orange")) +
  theme(
    panel.border = element_blank(),
    axis.text = element_blank(),
    axis.title.x=element_blank(),
    axis.title.y=element_blank(),
    panel.grid = element_blank(),
    axis.ticks = element_blank(),
    plot.background = element_rect(fill = "transparent"),
    panel.background = element_rect(fill = "transparent"),
    legend.position = ("right"),
    legend.title=element_blank())

#olfactory that were translocated 500
ggplot() + 
  geom_circle(aes(x0 = 0, y0 = 500, r = 200), linetype = "solid", color ="grey", fill = "grey", size = 0.25, alpha = 0.5) +
  geom_circle(aes(x0 = 0, y0 = 0, r = 100), linetype = "solid", size = 0.25, color = "grey") +
  geom_circle(aes(x0 = 0, y0 = 0, r = 200), linetype = "solid", size = 0.25, color = "grey") +
  geom_circle(aes(x0 = 0, y0 = 0, r = 300), linetype = "solid", size = 0.25, color = "grey") +
  geom_circle(aes(x0 = 0, y0 = 0, r = 400), linetype = "solid", size = 0.25, color = "grey") +
  geom_circle(aes(x0 = 0, y0 = 0, r = 500), linetype = "solid", size = 0.25, color = "grey") +
  geom_circle(aes(x0 = 0, y0 = 0, r = 600), linetype = "solid", size = 1, color = "black") +
  geom_segment(aes(x = -600, xend = 600, y = 0, yend = 0), linetype = "solid", size = 0.25, color = "grey") +
  geom_segment(aes(x = 0, xend = 0, y = -600, yend = 600), linetype = "solid", size = 0.25, color = "grey") + 
  geom_path(data = OA_df[(OA_df$Distance ==500 & OA_df$homing == 1),], aes(x = x_new, y = y_new, group = id, color = as.factor(Ablation)), arrow = arrow(angle = 20, length = unit(0.1, "inches"), ends = "last", type ="closed")) +
  geom_path(data = OA_df[(OA_df$Distance ==500 & OA_df$homing == 0),], aes(x = x_new, y = y_new, group = id, color = as.factor(Ablation)), arrow = arrow(angle = 20, length = unit(0.1, "inches"), ends = "last", type ="closed")) +
  coord_fixed(ratio = 1)+
  scale_color_manual(values = c("darkgreen", "orange")) +
  theme(
    panel.border = element_blank(),
    axis.text = element_blank(),
    axis.title.x=element_blank(),
    axis.title.y=element_blank(),
    panel.grid = element_blank(),
    axis.ticks = element_blank(),
    plot.background = element_rect(fill = "transparent"),
    panel.background = element_rect(fill = "transparent"),
    legend.position = "right",
    legend.title=element_blank())

#####Figure 3C - Magnetroeception
#megnetoreception that were translocated 1000
#remove MAE* toads
MA_df <- toads_df[toads_df$Ablation == "MAE" | toads_df$Ablation == "MAC",]

ggplot() + 
  geom_circle(aes(x0 = 0, y0 = 1000, r = 200), linetype = "solid", color ="grey", fill = "grey", size = 0.25, alpha = 0.5) +
  geom_circle(aes(x0 = 0, y0 = 0, r = 200), linetype = "solid", size = 0.25, color = "grey") +
  geom_circle(aes(x0 = 0, y0 = 0, r = 400), linetype = "solid", size = 0.25, color = "grey") +
  geom_circle(aes(x0 = 0, y0 = 0, r = 600), linetype = "solid", size = 0.25, color = "grey") +
  geom_circle(aes(x0 = 0, y0 = 0, r = 800), linetype = "solid", size = 0.25, color = "grey") +
  geom_circle(aes(x0 = 0, y0 = 0, r = 1000), linetype = "solid", size = 0.25, color = "grey") +
  geom_circle(aes(x0 = 0, y0 = 0, r = 1200), linetype = "solid", size = 1, color = "black") +
  geom_segment(aes(x = -1200, xend = 1200, y = 0, yend = 0), linetype = "solid", size = 0.25, color = "grey") +
  geom_segment(aes(x = 0, xend = 0, y = -1200, yend = 1200), linetype = "solid", size = 0.25, color = "grey") + 
  geom_path(data = MA_df[(MA_df$Distance ==1000 & MA_df$homing == 1),], aes(x = x_new, y = y_new, group = id, color = as.factor(Ablation)), arrow = arrow(angle = 20, length = unit(0.1, "inches"), ends = "last", type ="closed")) +
  geom_path(data = MA_df[(MA_df$Distance ==1000 & MA_df$homing == 0),], aes(x = x_new, y = y_new, group = id, color = as.factor(Ablation)), arrow = arrow(angle = 20, length = unit(0.1, "inches"), ends = "last", type ="closed")) +
  coord_fixed(ratio = 1)+
  scale_color_manual(values = c("yellow4", "violet")) +
  theme(
    panel.border = element_blank(),
    axis.text = element_blank(),
    axis.title.x=element_blank(),
    axis.title.y=element_blank(),
    panel.grid = element_blank(),
    axis.ticks = element_blank(),
    plot.background = element_rect(fill = "transparent"),
    panel.background = element_rect(fill = "transparent"),
    legend.position = ("right"),
    legend.title=element_blank())

#magnetoreception that were translocated 500
ggplot() + 
  geom_circle(aes(x0 = 0, y0 = 500, r = 200), linetype = "solid", color ="grey", fill = "grey", size = 0.25, alpha = 0.5) +
  geom_circle(aes(x0 = 0, y0 = 0, r = 100), linetype = "solid", size = 0.25, color = "grey") +
  geom_circle(aes(x0 = 0, y0 = 0, r = 200), linetype = "solid", size = 0.25, color = "grey") +
  geom_circle(aes(x0 = 0, y0 = 0, r = 300), linetype = "solid", size = 0.25, color = "grey") +
  geom_circle(aes(x0 = 0, y0 = 0, r = 400), linetype = "solid", size = 0.25, color = "grey") +
  geom_circle(aes(x0 = 0, y0 = 0, r = 500), linetype = "solid", size = 0.25, color = "grey") +
  geom_circle(aes(x0 = 0, y0 = 0, r = 600), linetype = "solid", size = 1, color = "black") +
  geom_segment(aes(x = -600, xend = 600, y = 0, yend = 0), linetype = "solid", size = 0.25, color = "grey") +
  geom_segment(aes(x = 0, xend = 0, y = -600, yend = 600), linetype = "solid", size = 0.25, color = "grey") + 
  geom_path(data = MA_df[(MA_df$Distance ==500 & MA_df$homing == 1),], aes(x = x_new, y = y_new, group = id, color = as.factor(Ablation)), arrow = arrow(angle = 20, length = unit(0.1, "inches"), ends = "last", type ="closed")) +
  geom_path(data = MA_df[(MA_df$Distance ==500 & MA_df$homing == 0),], aes(x = x_new, y = y_new, group = id, color = as.factor(Ablation)), arrow = arrow(angle = 20, length = unit(0.1, "inches"), ends = "last", type ="closed")) +
  coord_fixed(ratio = 1)+
  scale_color_manual(values = c("yellow4", "violet", "darkgray")) +
  theme(
    panel.border = element_blank(),
    axis.text = element_blank(),
    axis.title.x=element_blank(),
    axis.title.y=element_blank(),
    panel.grid = element_blank(),
    axis.ticks = element_blank(),
    plot.background = element_rect(fill = "transparent"),
    panel.background = element_rect(fill = "transparent"),
    legend.position = "right",
    legend.title=element_blank())
###########################################################################################################
#PART 6
##Plot Temporal aspects of trajectories
##########################################################################################################
#Figure 2C: All Toads 500 m
ggplot(toads_df[toads_df$Distance==500, ], aes(x = time_elapsed, y = home_dist, color = as.factor(homing))) + 
  geom_rect(aes(xmin = 0, xmax = 75, ymin = 0, ymax = 200), linetype = "solid", size = 0.25, color = "grey",fill = "grey", alpha = 0.1 )+
  geom_path(aes(group = id), arrow = arrow(angle = 30, length = unit(0.1, "inches"), ends = "last", type ="closed")) + 
  geom_point(size = 2, shape = 1) +
  geom_segment(aes(x = 0, xend = 75, y = 200, yend = 200), linetype = "dashed", size = 0.25, color = "black") +
  xlim(0,90)+
  scale_color_manual(values = c("goldenrod", "blue")) +
  theme_classic() +
  theme(legend.title=element_blank())+
  theme(aspect.ratio=1)+
  theme(axis.title.x = element_text(size = 24, face = "bold", family = "Arial"), # X axis title
        axis.title.y = element_text(size = 24, face = "bold", family = "Arial"), # Y axis title
        axis.text.x = element_text(size = 22, family = "Arial"), # X axis text (ticks)
        axis.text.y = element_text(size = 22, family = "Arial"))+ # Y axis text (ticks)
  coord_fixed(ratio = 0.1)+
  xlab("Time Elapsed (hrs)")+
  ylab("Distance from Home (m)")+
  scale_y_continuous(breaks=seq(0, 600, by = 100))+
  scale_x_continuous(breaks = seq(24, 72, by = 24))

#Figure 2D: All Toads 1000 m
ggplot(toads_df[toads_df$Distance==1000, ], aes(x = time_elapsed, y = home_dist, color = as.factor(homing))) + 
  geom_rect(aes(xmin = 0, xmax = 125, ymin = 0, ymax = 200), linetype = "solid", size = 0.25, color = "grey",fill = "grey", alpha = 0.1 )+
  geom_path(aes(group = id), arrow = arrow(angle = 30, length = unit(0.1, "inches"), ends = "last", type ="closed")) + 
  geom_point(size = 2, shape = 1) +
  geom_segment(aes(x = 0, xend = 125, y = 200, yend = 200), linetype = "dashed", size = 0.25, color = "black") +
  xlim(0,90)+
  scale_color_manual(values = c("goldenrod", "blue")) +
  theme_classic() +
  theme(legend.title=element_blank())+
  theme(aspect.ratio=1)+
  theme(axis.title.x = element_text(size = 24, face = "bold", family = "Arial"), # X axis title
        axis.title.y = element_text(size = 24, face = "bold", family = "Arial"), # Y axis title
        axis.text.x = element_text(size = 22, family = "Arial"), # X axis text (ticks)
        axis.text.y = element_text(size = 22, family = "Arial"))+ # Y axis text (ticks)
  coord_fixed(ratio = 0.1)+
  xlab("Time Elapsed (hrs)")+
  ylab("Distance from Home (m)")+
  scale_y_continuous(breaks=seq(0, 1600, by = 200))+
  scale_x_continuous(breaks = seq(24, 120, by = 24))
###########################################################################################################
#Part 7
#Visualizations

########################################################################################################
#FIGURE1 Movement range
#FIGURE 1A - movement by range by field site

ggplot(base_summary, aes(x = loc, y=base.range, fill = loc)) +   
  geom_boxplot(outlier.shape = NA) + 
  geom_point(aes(shape = sex),
             position=position_jitterdodge(dodge.width=0.9), size=6) +
  ylab("Range (meters)")+
  scale_fill_manual(values = loc_colors)+
  coord_fixed(ratio = 0.003)+
  scale_y_continuous(breaks=c(0, 100, 200, 300, 400, 500, 600, 700),
                     labels=c('0', '100', '200', '300', '400', '500', '600', '700'))+
  theme(
    axis.title.y = element_blank(), # Y axis title
    axis.text.x = element_blank(), # X axis text (ticks)
    axis.text.y = element_text(size = 45, family = "Arial"), # Y axis text (ticks)
    panel.background = element_rect(fill = "transparent"),
    axis.title.x=element_blank(),
    legend.position = "none",
    legend.title=element_blank())
########################################################################################################
#Figure 2 Homing
#Figures 2A and 2B are plotted in Normalize&PlotTraj_052924.R
#Figures 2C and 2D are plotted in TemporalPlots_052924.R

#FIGURE 2E- Homing duration by field site
ggplot(homing, aes(x = loc, y=duration, fill = loc)) + 
  geom_boxplot(outlier.shape = NA) + geom_jitter(shape=16, position=position_jitter(0.1), size = 5)+
  ylab("Homing Duration (hrs)")+
  scale_fill_manual(values = loc_colors)+
  coord_fixed(ratio = 0.04)+
  theme(
    axis.title.y = element_blank(), # Y axis title
    axis.text.x = element_blank(), # X axis text (ticks)
    axis.text.y = element_text(size = 40, family = "Arial"), # Y axis text (ticks)
    panel.background = element_rect(fill = "transparent"),
    axis.title.x=element_blank(),
    legend.position = "none",
    legend.title=element_blank())

#FIGURE 2F- Active homing duration by field site

ggplot(homing, aes(x = loc, y=homing_time, fill = loc)) + 
  geom_boxplot(outlier.shape = NA) + geom_jitter(shape=16, position=position_jitter(0.1), size = 5)+
  ylab("Homing Duration (hrs)")+
  scale_fill_manual(values = loc_colors)+
  coord_fixed(ratio = 0.04)+
  theme(
    axis.title.y = element_blank(), # Y axis title
    axis.text.x = element_blank(), # X axis text (ticks)
    axis.text.y = element_text(size = 40, family = "Arial"), # Y axis text (ticks)
    panel.background = element_rect(fill = "transparent"),
    axis.title.x=element_blank(),
    legend.position = "none",
    legend.title=element_blank())
########################################################################################################
#FIGURE 3 Ablation experiments
#Figure 3A is plotted in Normalize&PlotTraj_052924.R

#Figure 3B- Cumulative distance for olfaction ablated and sham animals
ggplot(traj_OA_summary, aes(x = as.factor(con), y= cumulative, fill = con)) + 
  geom_boxplot(outlier.shape = NA) + geom_jitter(shape=16, position=position_jitter(0.1), size = 5)+
  ylab("cumulative movement(m)")+
  xlab("ablation")+
  scale_fill_manual(values = ab_colors)+
  coord_fixed(ratio = 0.001)+
  theme(
    axis.title.y = element_blank(), # Y axis title
    axis.text.x = element_blank(), # X axis text (ticks)
    axis.text.y = element_text(size = 30, family = "Arial"), # Y axis text (ticks)
    panel.background = element_rect(fill = "transparent"),
    axis.title.x=element_blank(),
    legend.position = "none",
    legend.title=element_blank())

#Figure 3C is plotted in Normalize&PlotTraj_052924.R
#Figure 3D - cumulative movement in sham and experimental animals in magnetoreception experiment
ggplot(traj_MA_summary, aes(x = as.factor(con), y= cumulative, fill = con)) + 
  geom_boxplot(outlier.shape = NA) + geom_jitter(shape=16, position=position_jitter(0.1), size = 5)+
  ylab("cumulative movement(m)")+
  xlab("ablation")+
  scale_fill_manual(values = ab_colors)+
  coord_fixed(ratio = 0.001)+
  theme(
    axis.title.y = element_blank(), # Y axis title
    axis.text.x = element_blank(), # X axis text (ticks)
    axis.text.y = element_text(size = 30, family = "Arial"), # Y axis text (ticks)
    panel.background = element_rect(fill = "transparent"),
    axis.title.x=element_blank(),
    legend.position = "none",
    legend.title=element_blank())

########################################################################################################

#FIGURE 4 Brain activity
#Figure 4A - Brain activity comparison between all toads
ggboxplot(ps6_means, y ="density", x="Brain.Region", color="group",
          add = "jitter") +
  scale_color_manual(values = con_colors)+
  scale_x_discrete(limits = c("DP", "LP", "MP","Ms","Ls","Stri"))+
  theme(
    panel.border = element_blank(),
    axis.text = element_blank(),
    panel.grid = element_blank(),
    axis.ticks = element_blank(),
    plot.background = element_rect(fill = "transparent"),
    panel.background = element_rect(fill = "transparent"),
    legend.position = ("right"),
    legend.title=element_blank())+
  coord_fixed(ratio = 0.004)+
  ylab("pS6-positive cells/mm")
########################################################################################################

#SUPPLEMENTAL FIGURES

########################################################################################################

#Figure S5 - movement range plotted against svl
mS5 <- lm(base.range~svl, data = base_summary)
mS5.plot <- ggplot(base_summary, aes(x = svl, y = base.range, color = loc)) + 
  geom_jitter(width = 0, height = 0.05) + 
  coord_fixed(ratio = 0.01)+
  scale_color_manual(values = loc_colors) +
  theme(
    panel.grid = element_blank(),
    axis.line = element_line(size = 1, colour = "black"),
    plot.background = element_rect(fill = "transparent"),
    legend.position = "right",
    legend.title=element_blank())
plot(mS5.plot)

#make a list of potential svls
svls <- seq(8, 17, 1) # predict svls

#make a predicted line for the model
predicted.mS5 <- predict(mS5, newdata = data.frame(svl = svls),type = "response")

mS5.plot.lines <- mS5.plot + 
  geom_line(data = data.frame(svl = svls, base.range = predicted.mS5),color = "black", linewidth = 0.5)+
  geom_smooth(method = "lm", se = TRUE, color = "black")+
  geom_point(size = 5)+
  ylab("movement range (m)")+
  xlab("snout-vent-length (cm)")+
  theme(
    axis.text.x = element_text(size = 30, family = "Arial"), # X axis text (ticks)
    axis.text.y = element_text(size = 30, family = "Arial"), # Y axis text (ticks)
    axis.title.y = element_text(size = 40, family = "Arial"), # Y axis text (ticks)
    panel.background = element_rect(fill = "transparent"),
    axis.title.x=element_text(size = 40, family = "Arial"),
    legend.position = "none",
    legend.title=element_blank())
plot_grid(mS5.plot.lines)

########################################################################################################

#S6A-Comparing homing duration between 500 and 1000 meter translocations
S6A <- ggboxplot(homing, x = "Distance", y = "duration", fill = "Distance",
                 add = "jitter")+
  ylab("Homing Duration (hrs)")+
  xlab("Translocation Distance")+
  theme(
    axis.text.x = element_text(size = 20, family = "Arial"), # X axis text (ticks)
    axis.text.y = element_text(size = 20, family = "Arial"), # Y axis text (ticks)
    axis.title.y = element_text(size = 40, family = "Arial"), # Y axis text (ticks)
    panel.background = element_rect(fill = "transparent"),
    axis.title.x=element_text(size = 40, family = "Arial"),
    legend.position = "none",
    legend.title=element_blank())
S6A


#S6B-Comparing homing speed between 500 and 1000 meter translocations

S6B <- ggboxplot(homing, x = "Distance", y = "homing_speed", fill = "Distance",
                 add = "jitter")+
  ylab("Homing Speed (m/hr)")+
  xlab("Translocation Distance")+
  coord_fixed(ratio = 0.005)+
  theme(
    axis.text.x = element_text(size = 20, family = "Arial"), # X axis text (ticks)
    axis.text.y = element_text(size = 20, family = "Arial"), # Y axis text (ticks)
    axis.title.y = element_text(size = 40, family = "Arial"), # Y axis text (ticks)
    panel.background = element_rect(fill = "transparent"),
    axis.title.x=element_text(size = 40, family = "Arial"),
    legend.position = "none",
    legend.title=element_blank())
S6B

########################################################################################################
#Figure S7- comparing duration by size

#S7A - relationship between size and homing furation
mS7A <- lm(duration~svl, data = homing)
summary(mS7A)
mS7A.plot <- ggplot(homing, aes(x = svl, y = duration, color = loc)) + 
  geom_jitter(width = 0, height = 0.05) + 
  coord_fixed(ratio = 0.1)+
  scale_color_manual(values = loc_colors) +
  theme(
    panel.grid = element_blank(),
    axis.line = element_line(size = 1, colour = "black"),
    plot.background = element_rect(fill = "transparent"),
    legend.position = "right",
    legend.title=element_blank())
plot(mS7A.plot)

#make a list of potential svls
svls <- seq(8, 17, 1) # predict svls

#make a predicted line for the model
predicted.mS7A <- predict(mS7A, newdata = data.frame(svl = svls),type = "response")

mS7A.plot.lines <- mS7A.plot + 
  geom_line(data = data.frame(svl = svls, duration = predicted.mS7A),color = "black", linewidth = 0.5)+
  geom_smooth(method = "lm", se = TRUE, color = "black")+
  geom_point(size = 5)+
  ylab("duration (hrs)")+
  xlab("snout-vent-length (cm)")+
  theme(
    axis.text.x = element_text(size = 30, family = "Arial"), # X axis text (ticks)
    axis.text.y = element_text(size = 30, family = "Arial"), # Y axis text (ticks)
    axis.title.y = element_text(size = 40, family = "Arial"), # Y axis text (ticks)
    panel.background = element_rect(fill = "transparent"),
    axis.title.x=element_text(size = 40, family = "Arial"),
    legend.position = "none",
    legend.title=element_blank())
plot_grid(mS7A.plot.lines)

#S7B
#PLOT SVL by SITE

ggplot(traj_all_summary[traj_all_summary$homing == 1,], aes(x = loc, y=svl, fill = loc)) + 
  geom_boxplot(outlier.shape = NA) + geom_jitter(shape=16, position=position_jitter(0.1), size = 5)+
  ylab("snout-vent-length")+
  scale_fill_manual(values = loc_colors)+
  coord_fixed(ratio = 0.5)+
  theme(
    axis.title.y = element_blank(), # Y axis title
    axis.text.x = element_blank(), # X axis text (ticks)
    axis.text.y = element_text(size = 40, family = "Arial"), # Y axis text (ticks)
    panel.background = element_rect(fill = "transparent"),
    axis.title.x=element_blank(),
    legend.position = "none",
    legend.title=element_blank())

#normal
shapiro.test(homing$svl)
#ANOVA on only homing toads shown here
b<- aov(svl~loc, homing)
summary(b)
b_posthoc <- TukeyHSD(b)
b_posthoc

########################################################################################################
#FIGURE S8 Circular distributions of toad movements in relation to the home direction. 
#Plotted at the conclusion of HI_Toads_Spatial_Data_052924 in the circular stats section

########################################################################################################
#Figure S9 - ps6 active cells between sham and ablated animals
ggplot(olf_ct_means, aes(x = as.factor(Ablation), y= pS6, fill = Ablation)) + 
  geom_boxplot(outlier.shape = NA) + geom_jitter(shape=16, position=position_jitter(0.1), size = 5)+
  ylab("pS6 count")+
  xlab("ablation")+
  scale_fill_manual(values = ab_colors)+
  coord_fixed(ratio = 0.02)+
  theme(
    axis.title.y = element_blank(), # Y axis title
    axis.text.x = element_blank(), # X axis text (ticks)
    axis.text.y = element_text(size = 30, family = "Arial"), # Y axis text (ticks)
    panel.background = element_rect(fill = "transparent"),
    axis.title.x=element_blank(),
    legend.position = "none",
    legend.title=element_blank())
########################################################################################################

#FIGURE S10A Brain activity in Olfactary ablated and sham animals
ggboxplot(ps6_means2[ps6_means2$group == "OAC"| ps6_means2$group == "OAE",], y ="density", x="Brain.Region", color="group", palette = "jco",
          add = "jitter") +
  theme(
    panel.border = element_blank(),
    axis.text = element_blank(),
    panel.grid = element_blank(),
    axis.ticks = element_blank(),
    plot.background = element_rect(fill = "transparent"),
    panel.background = element_rect(fill = "transparent"),
    legend.position = ("right"),
    legend.title=element_blank())+
  coord_fixed(ratio = 0.003)+
  scale_color_manual(values = ab_colors)+
  scale_x_discrete(limits = c("DP", "LP", "MP","Ms","Ls","Stri"))+
  ylab("pS6-positive cells/mm") 

#FIGURE S10B Brain activity in Magnetoreception ablated and sham animals
ggboxplot(ps6_means2[ps6_means2$group == "MAC"| ps6_means2$group == "MAE",], y ="density", x="Brain.Region", color="group", palette = "jco",
          add = "jitter") +
  theme(
    panel.border = element_blank(),
    axis.text = element_blank(),
    panel.grid = element_blank(),
    axis.ticks = element_blank(),
    plot.background = element_rect(fill = "transparent"),
    panel.background = element_rect(fill = "transparent"),
    legend.position = ("right"),
    legend.title=element_blank())+
  coord_fixed(ratio = 0.003)+
  scale_x_discrete(limits = c("DP", "LP", "MP","Ms","Ls","Stri"))+
  scale_color_manual(values = ab_colors)+
  ylab("pS6-positive cells/mm") 
```
