## Supplementary material for "Neural and sensory basis of homing behavior in the invasive cane toad, *Rhinella marina*": R Scripts for Analysis: READ_ME_code.rtf

In this study, we characterize homing behavior In invasive cane toads in Hawaii, test the effects of sensory manipulations on toad navigation, and characterize brain activity differences between homing and non-homing toads.

The attached R script contains all code for analysis in the manuscript. All necessary packages are listed in the beginning of the code. The script references csv datasheets that have been uploaded as supplementary material in an Excel workbook and as csv files. The script is split into 7 sections:

PART 1: Analysis of translocation related spatial activity (lines 71-612)
PART 2: Analysis of Baseline Tracking Data (lines 613-760)
PART 3: Script for looking at effects of ablations on pre-translocation movements (lines 761-923)
PART 4: Analysis of pS6 count data (lines 924-1212)
PART 5: Normalize trajectories and plot with origin being translocation release site (lines 1213-1408)
PART 6: Plot Temporal aspects of trajectories (lines 1409-1453)
PART 7: Visualizations (lines 1454-1787)

In order to run this code, you must download the provided datafile and change the working directories and file paths throughout the code to the location where you are storing the data. 
