## Supplementary material for "Neural and sensory basis of homing behavior in the invasive cane toad, *Rhinella marina*": Datasheets: READ_ME_data.rtf

In this study, we characterize homing behavior In invasive cane toads in Hawaii, test the effects of sensory manipulations on toad navigation, and characterize brain activity differences between homing and non-homing toads.

In this excel workbook and CSV files, you will find all the datasets associated with the manuscript. The first sheet contains a landing page with descriptions of all datasets, and links to the other sheets where the data is recorded. These sheets correspond to the CSV files references in the R code also submitted in the supplemental material.

There are 6 sheets containing datasets related to spatial, behavioral component of the paper:
Baseline
Ablated_Baseline
Translocation_Coordinates
Transect_Correction
HI 2022 Toad List
TansToadInfo

There are 4 sheets containing datasets related to the analysis of neural activity:
Brains
Toads
BrainCounts_02
olf_ct

And there is one additional sheet which contains weather data from NOAA for each field site for the duration of translocation experiments:
Weather
