## Supplementary Figures, Tables, and Methods for "Neural and sensory basis of homing behavior in the invasive cane toad, *Rhinella marina*"

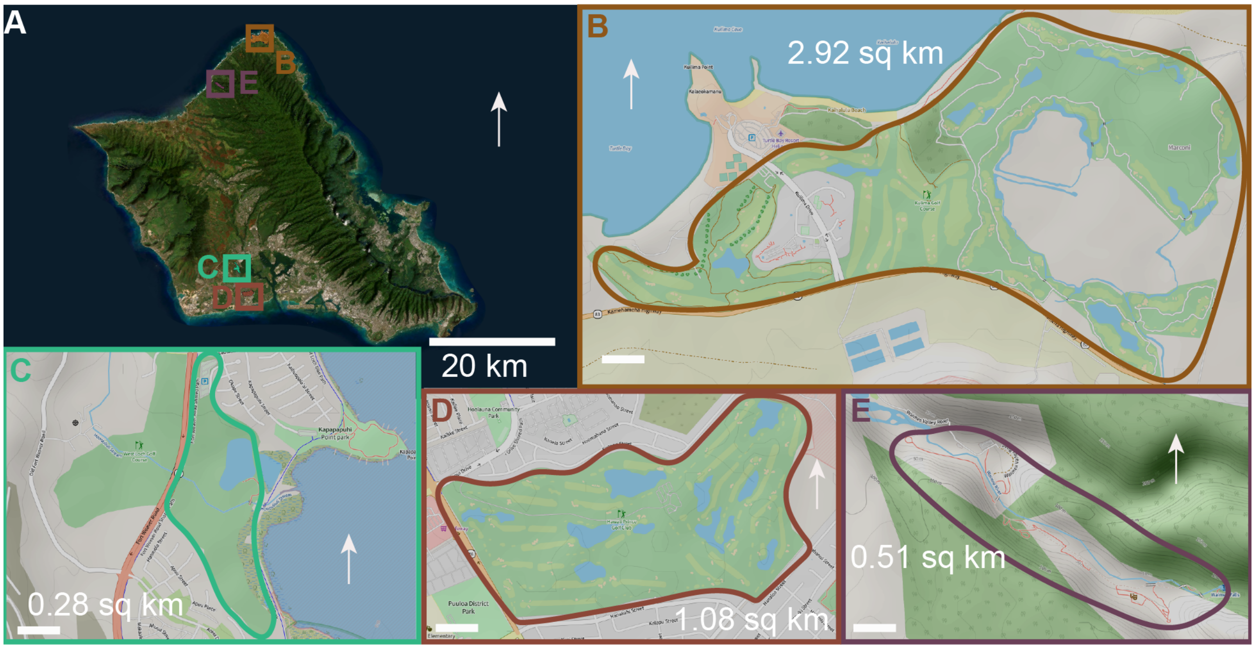


**Figure S1. Field sites on the island of Oahu**. White arrows point north. **(A)** Oahu with field sites labelled with corresponding panels. Imagery from ArcGis Online. **(B-E)** Each field site with tracking area outlined and approximate surface area provided in square kilometers. Images from OpenStreet Maps. Scale bars represent 200-m. **(B)** Turtle Bay (21° 42' 07.5" N 157° 59' 44.6” W), **(C)** West Loch (21° 22' 04.7" N 158° 01' 31.5" W), **(D)** Hawaii Prince (21° 18' 41.5" N 158° 00' 32.7" W), and **(E)** Waimea Valley (21° 38' 04.4" N 158° 03' 14.6” W).

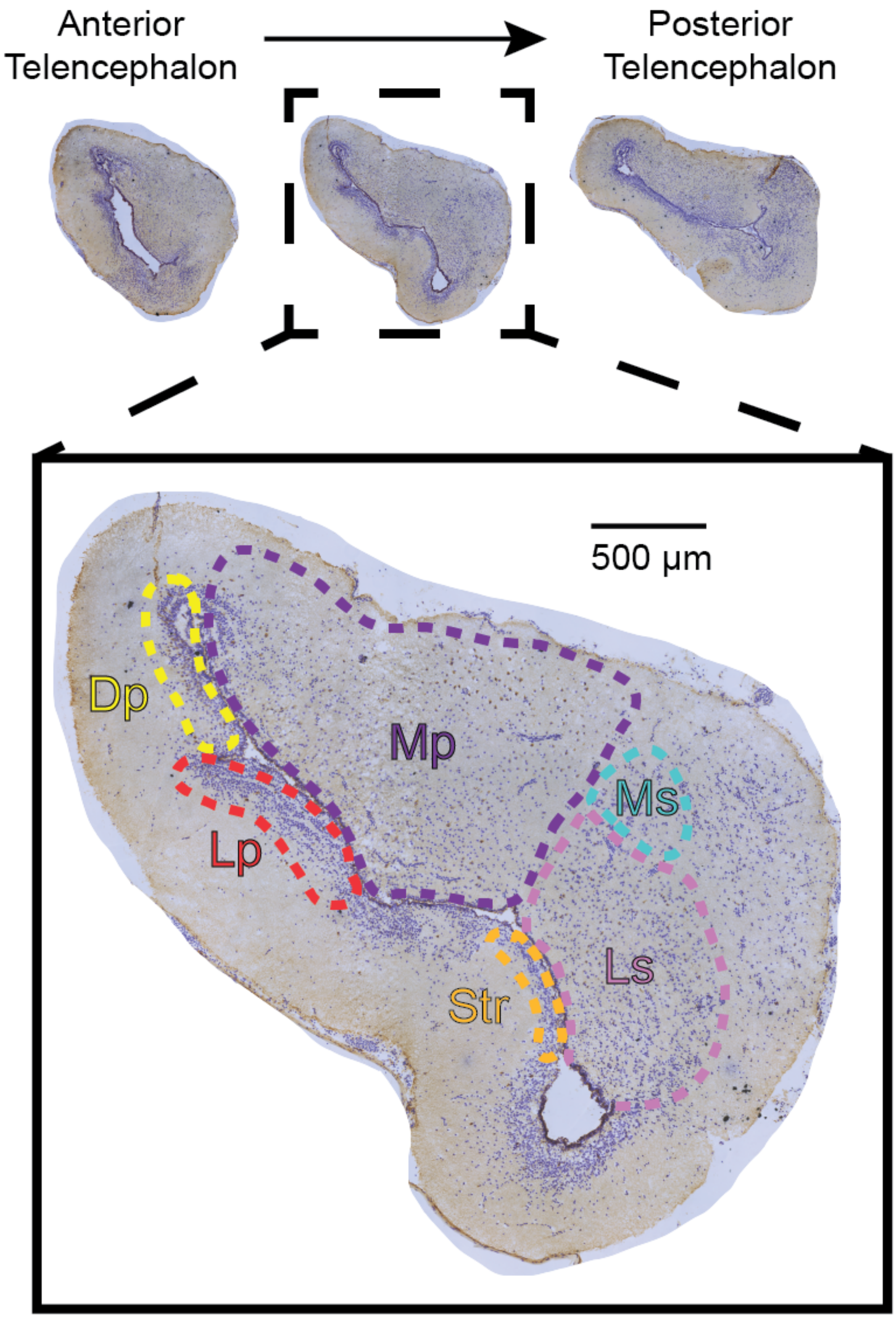


**Figure S2. Brain regions analyzed for activity.** Top of the image shows the anterior, middle, and posterior sections of the telencephalon. Three sections of each were counted for pS6-positive cells. Inset shows the location of the brain regions in the mid-telencephalic region. Abbreviations: Dp, dorsal pallium; Lp, lateral pallium; Mp, medial pallium; Ms, medial septum; Ls, lateral septum Str, striatum.

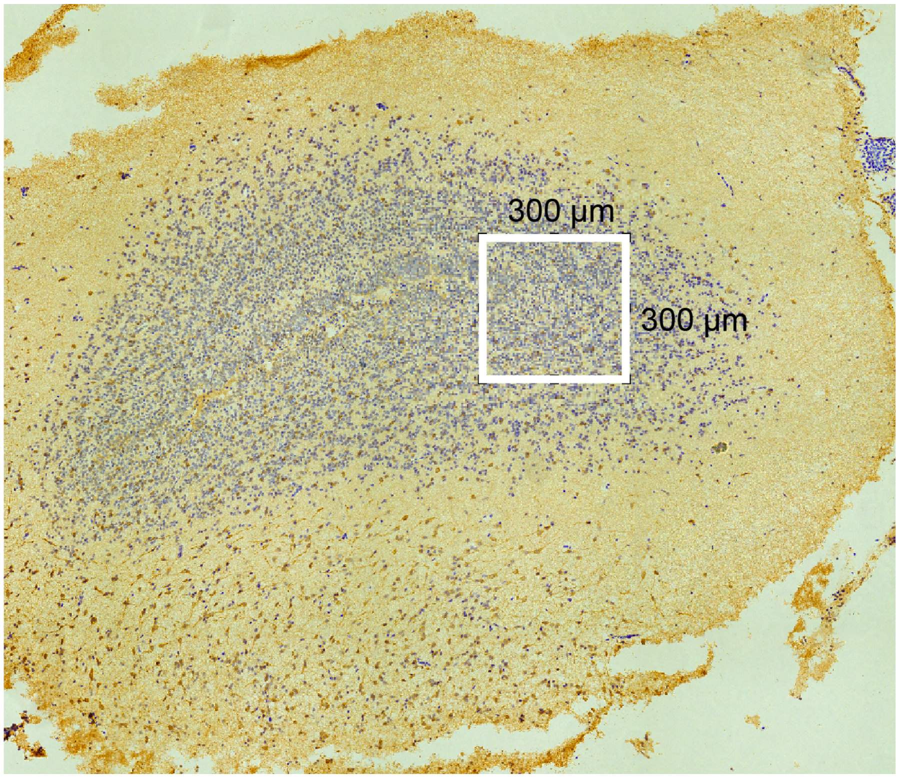


**Figure S3. Olfactory bulb cell quantification area.** Left Hemisphere of the olfactory bulb with 300 µm x 300 µm bounding box at the lateral boundary of the granule cell layer indicating where cells were counted within toads in the olfaction ablation experiment.

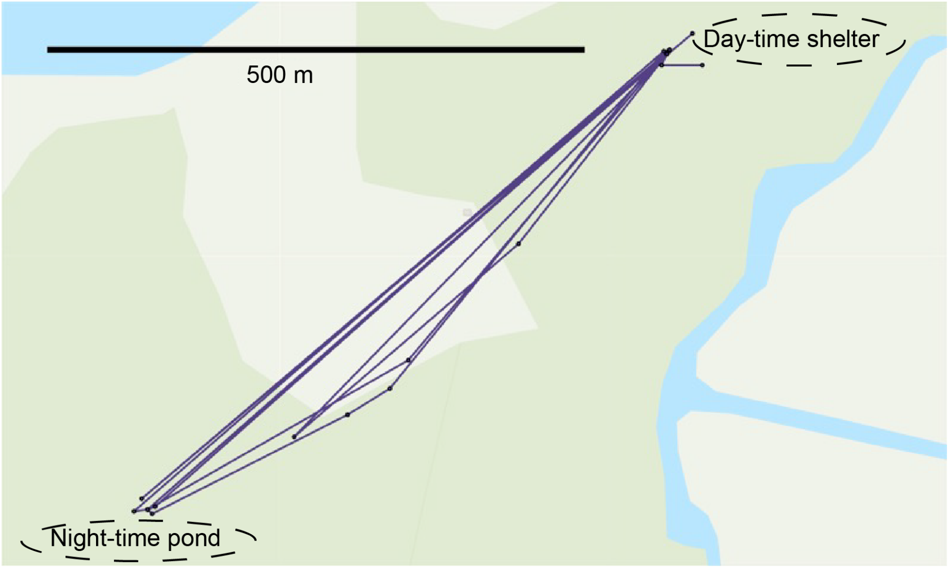


**Figure S4. Example of a "commuter toad."** Toad D6 made nightly movements between two ponds used as a night-time activity area and a day-time shelter.

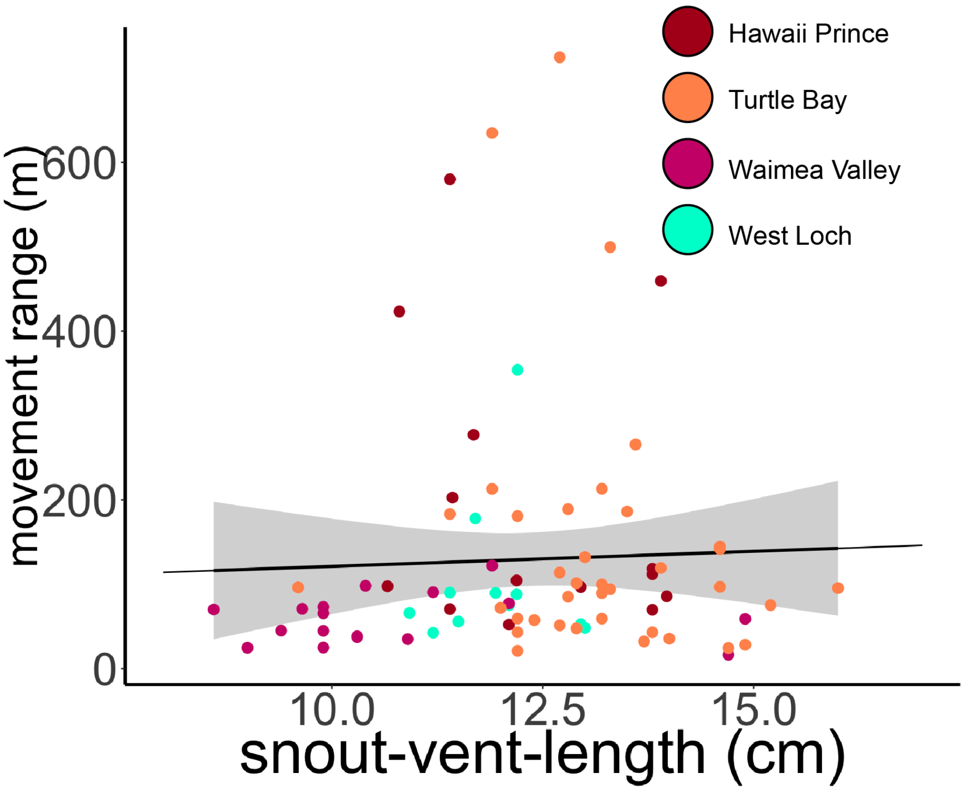


**Figure S5. Size does not affect movement range.** There was no significant effect of size (snout-vent-length) on toad movement range (linear mixed effects model: $t_{(68.60)} = -0.814$, $p = 0.42$). Gray Shading represents standard error.

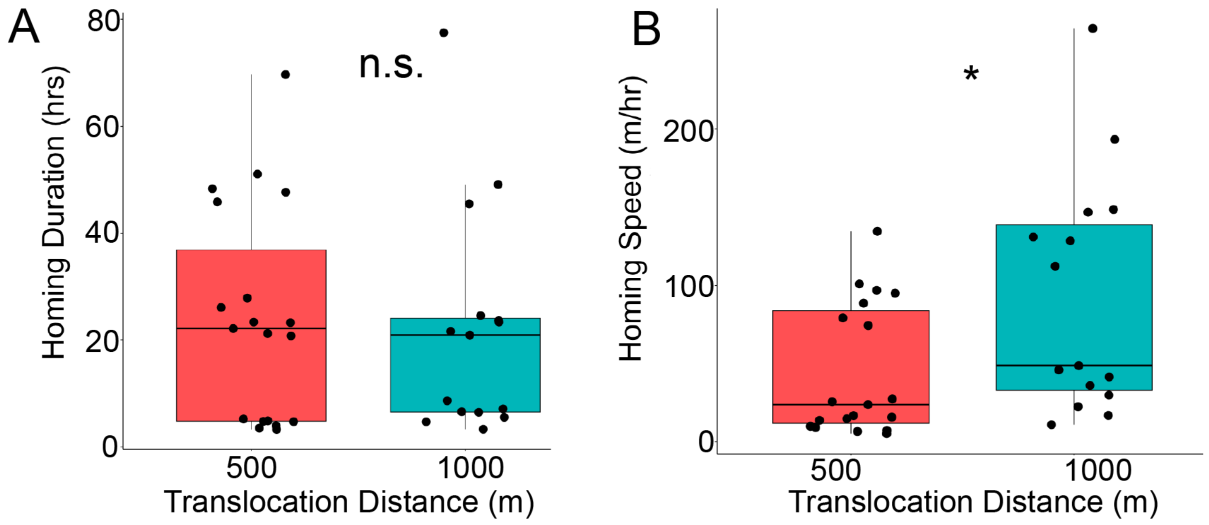


**Figure S6. Effects of translocation distance on homing. (A)** There was no significant effect of translocation distance on homing duration (Wilcoxon, W = 143.5, $p = 0.99$) because **(B)** toads homing from 1000 m moved faster (Wilcoxon, W = 72, $p = 0.015$).

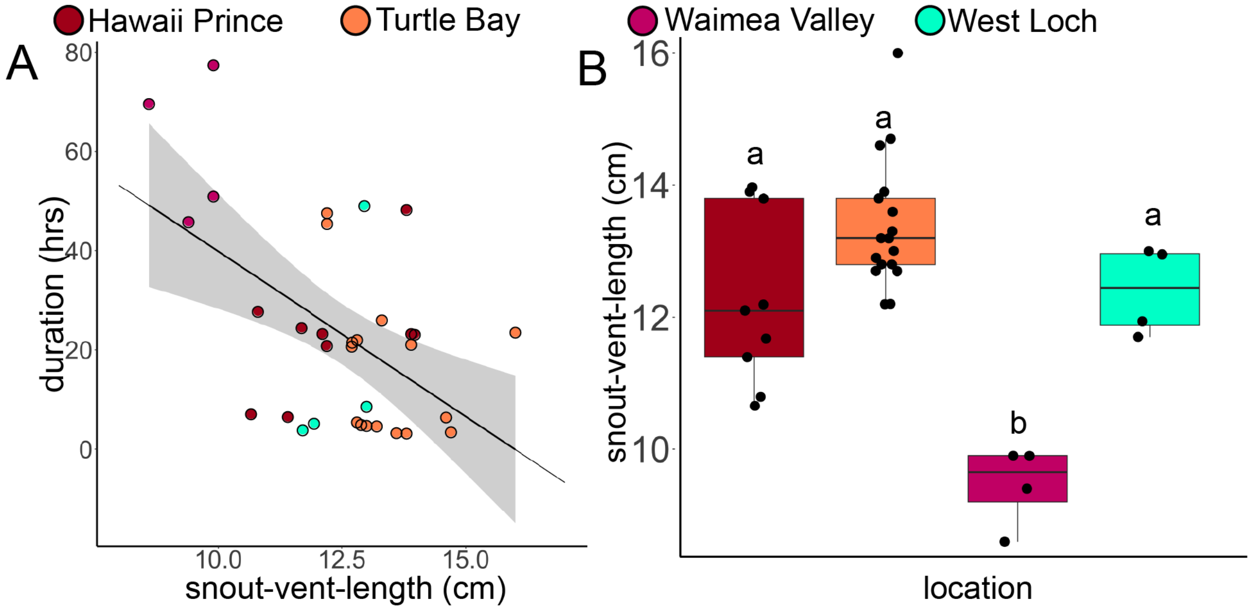


**Figure S7. Body size, homing speed, and study site relationship. (A)** While a linear model suggests toads with longer snout-vent-lengths returned home faster ($F_{(1,32)}$ = 12.03, p = 0.0015), the relationship is not significant when location is considered as a random effect ($t_{(30.52)}$ = -0.51, p = 0.61). Gray shading represents standard error. **(B)** Size of homing toads differed between field sites (ANOVA, $F_{(3)}$ = 16.12 , p = 1.99e-06) with Waimea Valley toads being the smallest and the slowest. Letters represent significance from post-hoc Tukey's test.

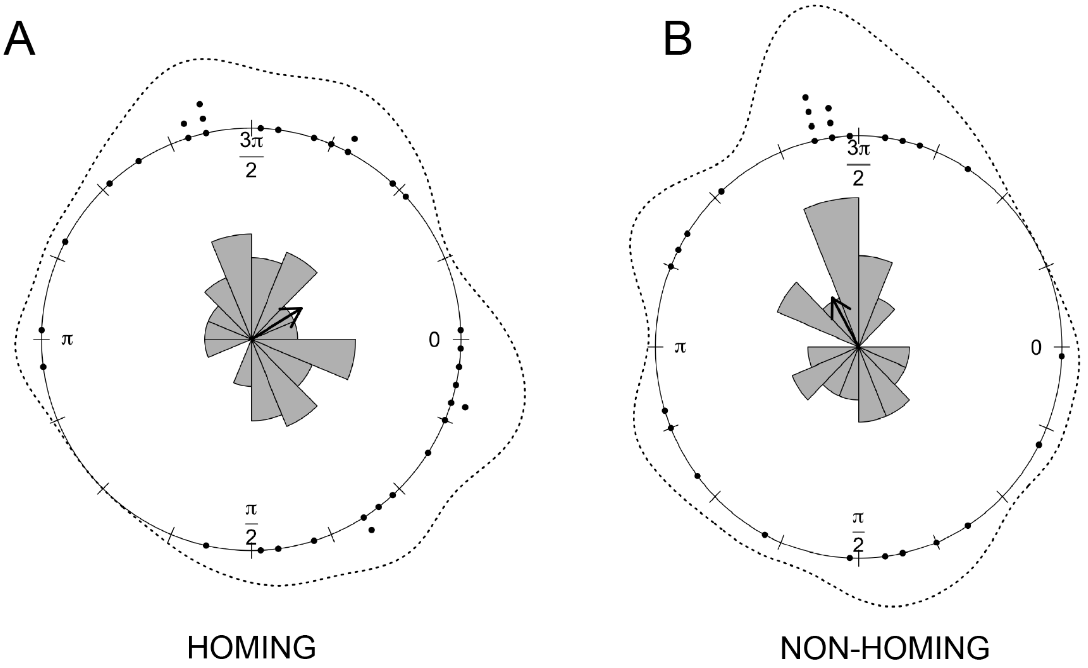


**Figure S8. Circular distributions of toad movements in relation to the home direction. (A)** The mean direction of travel of homing toads was significantly oriented towards home , shown as 0 in the plot (Rayleigh p-value = 0.024). **(B)** Direction of movement in non-homing toads was not significantly homewards oriented (Rayleigh p-value = 0.83). Individual dots represent the mean angle of movement for individual toads in relation to their home area. The dotted line surrounding the circle and wedges represent the distribution of these angles. Arrow inside of the plot indicates the mean angle and vector length of movement for all toads in that group.

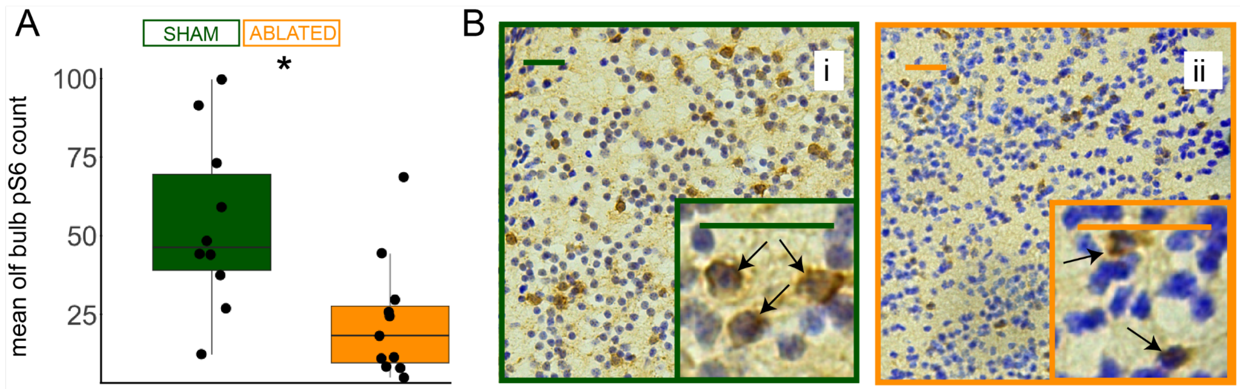


**Figure S9. Olfactory ablation leads to fewer active neurons in the olfactory bulb. (A)** Sham toads exhibited more pS6-positive cells in the olfactory bulb than ablated animals (Wilcoxon, W = 93.5, p = 0.0074). Each point represents the mean pS6 count across three sections of olfactory bulb in the same toad. **(B)** Representative areas of olfactory bulb (i) sham and (ii) ablated animals. Scale bars represent 25 μm. In inset, arrows indicate pS6-positive cells.

**Table S1. Summary of main statistical effects from generalized linear mixed model for differences in brain activity**

| | $\chi^2$ | df | p |
|---|---|---|---|
| Homing Condition | 14.085 | 2 | 0.0008738 |
| Brain Region | 2221.31 | 5 | < 2.2e-16 |
| Homing Condition x Brain Region | 112.800 | 10 | <2.2e-16 |

**Table S2. Pairwise contrasts by Homing Condition in medial pallium across AP axis**

| Section | Brain Region | Contrast | estimate | std. error | z-ratio | p-value |
|---|---|---|---|---|---|---|
| Anterior | Mp | Baseline-Homing | -0.6846 | 0.235 | -2.91 | 0.0101 |
| Anterior | Mp | Baseline-Nonhoming | -0.091 | 0.238 | -0.383 | 0.9224 |
| Anterior | Mp | Homing-Nonhoming | 0.5936 | 0.177 | 3.344 | 0.0024 |
| Medial | Mp | Baseline-Homing | -0.65462 | 0.207 | -3.155 | 0.0046 |
| Medial | Mp | Baseline-Nonhoming | -0.11899 | 0.21 | -0.567 | 0.8375 |
| Medial | Mp | Homing-Nonhoming | 0.53564 | 0.157 | 3.422 | 0.0018 |
| Posterior | Mp | Baseline-Homing | -0.7668 | 0.214 | -3.59 | 0.001 |
| Posterior | Mp | Baseline-Nonhoming | -0.0799 | 0.216 | -0.37 | 0.9272 |
| Posterior | Mp | Homing-Nonhoming | 0.6869 | 0.161 | 4.261 | 0.0001 |

*Shaded rows represent significant relationships

**Table S3. Mixed-effects linear modeling of brain activity as a function of path straightness and brain region**

| Fixed effects | Estimate | std. error | df | t-statistic | Pr(>\|t\|) |
|---|---|---|---|---|---|
| (Intercept) | 100.4485 | 61.583 | 39.4547 | 1.631 | 0.1108 |
| straightness | -1.8823 | 75.8438 | 39.4616 | -0.025 | 0.9803 |
| Brain.RegionLP | 0.3207 | 35.7288 | 1182.9999 | 0.009 | 0.9928 |
| Brain.RegionLs | 21.0416 | 39.1427 | 1183.0002 | 0.538 | 0.591 |
| Brain.RegionMP | 228.9358 | 35.7288 | 1182.9999 | 6.408 | 2.13E-10 |
| Brain.RegionMs | -56.7626 | 39.1427 | 1183.0002 | -1.45 | 0.1473 |
| Brain.RegionStr | -96.9117 | 39.1377 | 1182.9998 | -2.476 | 0.0134 |
| straightness:Brain.RegionLP | 44.0071 | 44.0091 | 1182.9998 | 1 | 0.3175 |
| straightness:Brain.RegionLs | 31.0172 | 48.21 | 1182.9999 | 0.643 | 0.5201 |
| straightness:Brain.RegionMP | -8.2816 | 44.0091 | 1182.9998 | -0.188 | 0.8508 |
| straightness:Brain.RegionMs | 40.9 | 48.21 | 1182.9999 | 0.848 | 0.3964 |
| straightness:Brain.RegionStr | 16.276 | 48.2094 | 1182.9998 | 0.338 | 0.7357 |

*Shaded rows represent significant relationships

**Table S4. Summary of individual mixed-effects linear models of brain activity as a function of path straightness for each brain region**

| Brain Region | Fixed effects | Estimate | Std. Error | df | t-value | Pr(>\|t\|) |
|---|---|---|---|---|---|---|
| | (Intercept) | 88.077 | 34.892 | 27.002 | 2.524 | 0.0178 |
| Dp | straightness | -1.758 | 42.971 | 27.003 | -0.041 | 0.9677 |
| | (Intercept) | 88.06 | 65.11 | 27 | 1.352 | 0.187 |
| Lp | straightness | 42.23 | 80.19 | 27 | 0.527 | 0.603 |
| | (Intercept) | 316.16 | 129.27 | 27.01 | 2.446 | 0.0212 |
| Mp | straightness | -10.6 | 159.19 | 27.01 | -0.067 | 0.9474 |
| | (Intercept) | 31.35 | 37.09 | 27.00 | 0.845 | 0.405 |
| Ms | straightness | 39.17 | 45.68 | 27.00 | 0.857 | 0.399 |
| | (Intercept) | 108.51 | 66.17 | 27.00 | 1.640 | 0.113 |
| Ls | straightness | 29.15 | 81.49 | 27.00 | 0.358 | 0.723 |
| | (Intercept) | -8.592 | 6.696 | 26.998 | -1.283 | 0.2103 |
| Str | straightness | 14.82 | 8.246 | 27 | 1.797 | 0.0835 |

**Table S5. Mixed-effects linear modeling of brain activity as a function of path duration**

| Fixed effects | Estimate | std. error | df | t-statistic | Pr(>\|t\|) |
|---|---|---|---|---|---|
| (Intercept) | 95.8429 | 18.3862 | 39.2191 | 5.213 | 6.30E-06 |
| duration | -0.4318 | 0.6651 | 39.218 | -0.649 | 0.51996 |
| Brain.RegionLP | 38.8002 | 10.5907 | 1182.9994 | 3.664 | 0.00026 |
| Brain.RegionLs | 74.062 | 11.6033 | 1183 | 6.383 | 2.49E-10 |
| Brain.RegionMP | 199.1365 | 10.5907 | 1182.9994 | 18.803 | < 2.00E-16 |
| Brain.RegionMs | -13.905 | 11.6033 | 1183 | -1.198 | 0.23101 |
| Brain.RegionStr | -89.842 | 11.6009 | 1182.9994 | -7.744 | 2.05E-14 |
| duration:Brain.RegionLP | -0.1895 | 0.3831 | 1182.9994 | -0.495 | 0.62098 |
| duration:Brain.RegionLs | -1.4169 | 0.4197 | 1183 | -3.376 | 0.00076 |
| duration:Brain.RegionMP | 1.0238 | 0.3831 | 1182.9994 | 2.673 | 0.00763 |
| duration:Brain.RegionMs | -0.5084 | 0.4197 | 1183 | -1.211 | 0.22602 |
| duration:Brain.RegionStr | 0.3108 | 0.4196 | 1182.9994 | 0.741 | 0.45901 |

*Shaded rows represent significant relationships

**Table S6. Summary of individual mixed-effects linear models of brain activity as a function of path duration for each brain region**

| Brain Region | Fixed effects | Estimate | Std. Error | df | t-value | Pr(>\|t\|) |
|---|---|---|---|---|---|---|
| | (Intercept) | 95.7076 | 10.2817 | 26.9994 | 9.309 | 6.46E-10 |
| Dp | duration | -0.4369 | 0.3719 | 26.9992 | -1.175 | 0.25 |
| | (Intercept) | 134.5563 | 19.4904 | 26.9987 | 6.904 | 2.03E-07 |
| Lp | duration | -0.6245 | 0.705 | 26.9986 | -0.886 | 0.384 |
| | (Intercept) | 295.271 | 38.923 | 27.015 | 7.586 | 3.67E-08 |
| Mp | duration | 0.603 | 1.408 | 27.015 | 0.428 | 0.672 |
| | (Intercept) | 81.9379 | 10.1958 | 27.0000 | 8.036 | 1.23E-08 |
| Ms | duration | -0.9402 | 0.3688 | 27.000 | -2.549 | 0.0168 |
| | (Intercept) | 169.9049 | 17.4584 | 27.0000 | 9.732 | 2.53E-10 |
| Ls | duration | -1.8486 | 0.6315 | 27.0000 | -2.927 | 0.00686 |
| | (Intercept) | 5.83011 | 2.03007 | 26.99628 | 2.872 | 0.00785 |
| Str | duration | -0.12742 | 0.07343 | 26.99596 | -1.735 | 0.0941 |

*Shaded rows represent significant relationships

**Table S7. Summary of main statistical effects for brain activity modeling in with ablation as a predictor**

| | $\chi^2$ | df | p-value |
|---|---|---|---|
| Homing Condition | 9.3652 | 1 | 0.002211 |
| Brain Region | 1996.0521 | 5 | < 2.20E-16 |
| Ablation | 1.4208 | 4 | 0.840571 |
| Homing Condition x Brain Region | 119.0922 | 5 | < 2.20E-16 |
| Condition x Ablation | 2.2813 | 4 | 0.684177 |
| Brain Region x Ablation | 127.5871 | 20 | < 2.20E-16 |
| Condition x Brain Region x Ablation | 155.5477 | 20 | < 2.20E-16 |

*Shaded rows represent significant effects

**Table S8. Pairwise contrasts by ablation condition in olfaction and magnetoreception ablation experiment**

| Brain Region | Contrast | estimate | std. error | z-ratio | p-value |
|---|---|---|---|---|---|
| Dp | OSH-OAB | 0.01817 | 0.267 | 0.068 | 1.0000 |
| Lp | OSH-OAB | -0.05879 | 0.262 | -0.224 | 0.9994 |
| Mp | OSH-OAB | -0.31512 | 0.262 | -1.205 | 0.7486 |
| Ms | OSH-OAB | 0.12493 | 0.267 | 0.467 | 0.9902 |
| Ls | OSH-OAB | 0.09948 | 0.266 | 0.374 | 0.9959 |
| Str | OSH-OAB | 0.05679 | 0.272 | 0.209 | 0.9996 |
| Dp | MSH-MAB | 0.17025 | 0.282 | 0.603 | 0.9746 |
| Lp | MSH-MAB | 0.10096 | 0.277 | 0.364 | 0.9963 |
| Mp | MSH-MAB | 0.22964 | 0.277 | 0.830 | 0.9215 |
| Ms | MSH-MAB | -0.27932 | 0.284 | -0.983 | 0.8629 |
| Ls | MSH-MAB | 0.15733 | 0.283 | 0.557 | 0.9811 |
| Str | MSH-MAB | -0.21646 | 0.290 | -0.747 | 0.9452 |

OSH = Olfactory Sham, OAB = Olfactory Ablated, MSH = Magnetoreception Sham, MAB = Magnetoreception Ablated

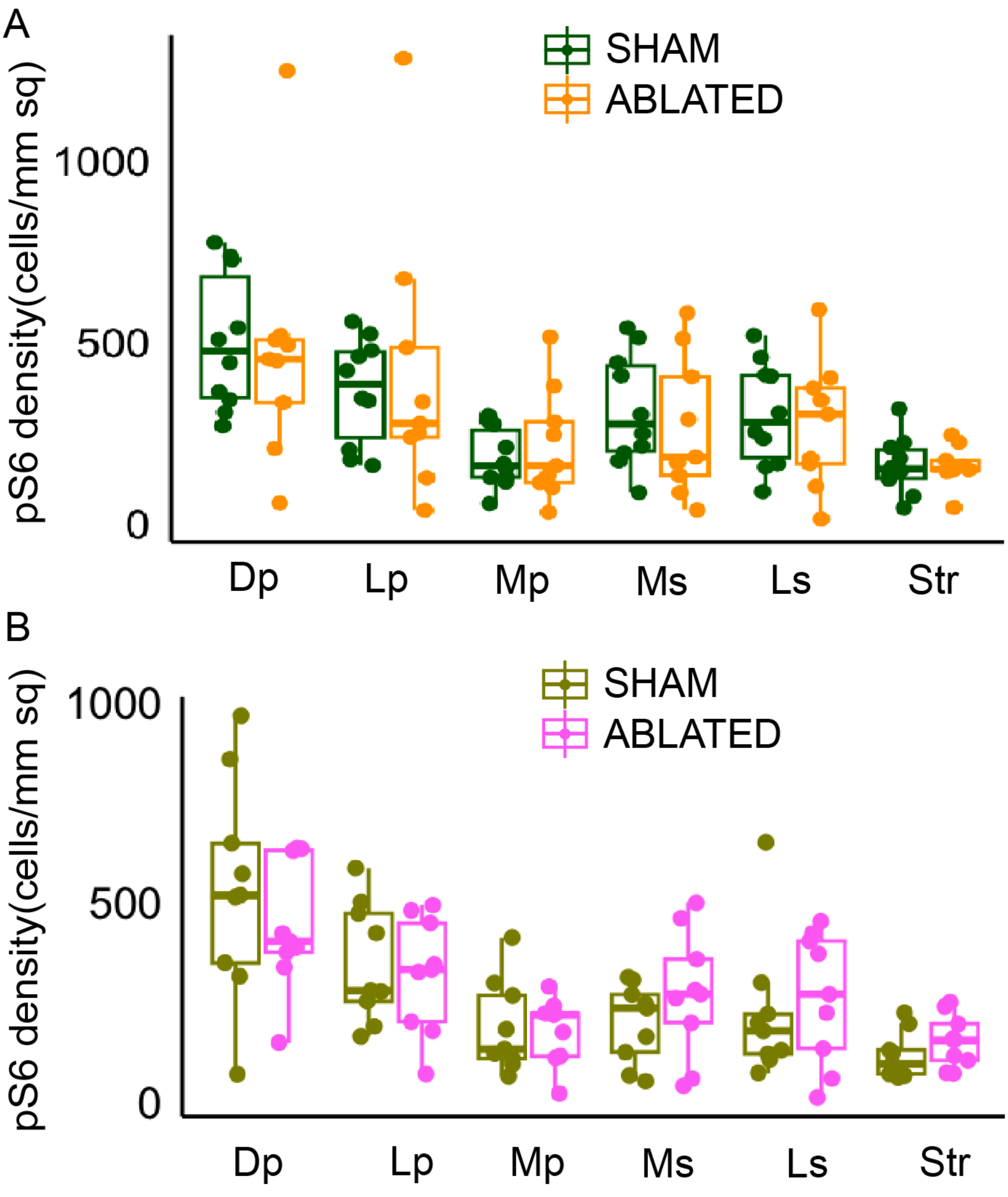


**Figure S10. Brain activity related to ablation in olfaction-ablated toads.** There were no significant pairwise differences in brain activity due to (**A**) olfaction or **(B)** magnetoreception ablation. Dp, dorsal pallium; Lp, lateral pallium; Mp, medial pallium; Sep, septum; Str, striatum.

**SM1. Ablation procedures**

In the olfactory ablation group (n = 11; 5 females and 6 males), toads were captured and lightly anesthetized through immersion in an MS-222 bath (1.4 g/L) for ~5 min. We performed olfactory ablations through zinc sulfate (ZnSO4) treatment, which temporarily damages the olfactory epithelium in juvenile and adult amphibians [1,2] and other vertebrates [3,4]. Once animals were anesthetized and not responding to toe pinches, they were held upside-down and each nostril was washed with 1% ZnSO4 dissolved in amphibian ringer solution (NaCl, KCl, $CaCl_2 \cdot 6H_2O$ and distilled water; Electron Microscopy Services, Hatfield, PA 19440) applied in five 200 µl doses through the internal nare with a P200 pipette. Any ZnSO4 that spilled into the animal's mouth was sucked up with the pipette. Following administration of ZnSO4, the toad's mouth was washed several times with amphibian ringer solution. Animals recovered in a net cage until fully mobile (usually 20-30 minutes), after which they were released at the capture point. Animals experiencing sham ablations (n = 11; 8 females and 3 males) underwent an identical procedure, but their nostrils were washed with amphibian ringer solution without ZnSO4.

Magnetoreception ablations were carried out by attaching a small Nickel plated Neodymium magnet (N50, 10 mm x 6 mm x 2 mm, ~1 g; Super Magnet Man, Pelham, Alabama, USA) with cyanoacrylate based adhesive (EM-150, Starbond, Torrance, CA, USA) to the skin over the skulls of toads (n = 11; 6 females and 5 males), similarly to previous studies in anurans [5–7]. Toads were captured, magnets applied, and toads were kept in a net cage for ~30 minutes before release. Sham ablation animals had a brass rectangle (12.7 mm x 6 mm x 1.6 mm, ~ 1 g; Metals Depot, Winchester, Kentucky, USA) attached in place of a magnet (n = 10; 3 females and 7 males). Two animals lost their magnets following translocation, so their behavior and brain measurements were not included in the magnetoreception analysis. Following translocation, animals were euthanized and tissues collected at timepoints described above (once toads either reached a point within 200 m of a baseline point or three days had elapsed post-translocation).

**SM2. pS6 immunohistochemistry with DAB visualization of a biotinylated antibody and cresyl violet staining for Nissl bodies**

*Remove slides from the -80°C freezer the day before staining begins

Day 1

1. Apply hydrophobic barrier around the edge of the slides with ImmEdge Hydrophobic Barrier PAP Pen (Vector Laboratories) and allow to dry for 3 minutes
2. Wash slides 3x for 10 min in 1x Tris-Buffered Saline (TBS)
3. Prepare Quenching Solution (~300 ul per slide)
    a. 3% of 30% $H_2O_2$
    b. 0.3% of TritonX-100
    c. 96.7% of 1x TBS
4. Apply Quenching Solution For 20 minutes
5. Prepare Blocking (~300 ul per slide)
    a. 5% Normal Goat Serum
    b. 0.3% TritonX-100
    c. 94.7% 1x TBS
6. Wash slides 3x for 10 min in 1x TBS
7. Apply blocking solution on slides for 1 hr at room temperature. Place slides onto glass trays in a closed baking tray with a small amount of water on the bottoms.
8. Ten min out from end of blocking incubation, prepare the primary antibody mix (~300 ul per slide)
    a. 1:300 pS6 primary (Phospho-S6 (Ser244, Ser247) Polyclonal Antibody, Invitrogen, Waltham, MA, USA)
    b. 2% Normal Goat Serum)
    c. 0.3% TritonX-100
    d. To volume with 1x TBS
9. Prepare identical mix without antibody for negative slides
10. Dump blocking solution of slides off and apply primary antibody mix. Incubate overnight in closed baking tray at room temperature

Day 2

1. Dump off the primary solution. Wash slides 3x for 10 min in 1x TBS
2. Make secondary antibody mix (~300 ul per slide)
    a. 1:200 biotinylated antibody (Goat Anti-Rat IgG Antibody (H+L), Vector Laboratories)
    b. 2% Normal Goat Serum
    c. 0.3% TritonX-100
    d. To volume with 1x TBS

3. Apply secondary mix to slides, and incubate for 2 hours in baking trays at room temperature.
4. Prepare DAB Solution
    a. 1% DAB (20x) in Distilled Water:
        i. Add 0.05g (50 mg) of DAB (3,3'-diaminobenzidine, Sigma Cat #D8001 or DAB-tetrahydrocholoride) to 5 ml distilled water. Add 10N HCl 2-3 drops (1 drop = 50 ul) and the solution turns light brown color. Shake for 10 minutes and DAB should dissolve completely. Keep in 4 °C until needed
    b. 0.3% $H_2O_2$ (20x) in distilled water:
        i. Add 50ul of 30% $H_2O_2$ in 5 ml distilled water and mix well. Store at 4 °C
5. Wash slides 3x for 10 min in 1x TBS. Make Avidin Biotin Complex (ABC) Horseradish Peroxidase (HRP) treatment (Vector Laboratories) by mixing 2 drops of each reagent from kit per 5 mL 1x TBS, ~300 ul needed per slide. ABC solution should stand for 15-30 min at RT before being applied to slides.
6. Apply ABC, incubate for 1 hr at room temperature in baking tray.
7. Wash slides 2x for 10 min in 1x TBS; then place slides into a large rack in 1x TBS in
8. Mix prepared DAB and $H_2O_2$ solutions in 500 mL 1x TBS, pH Adjust to 7.2, and filter. Put DAB solution into large dish
9. Put slides into DAB for 3.5 min or until brown color begins to develop on tissue.
10. End DAB reaction by moving slides into DI $H_2O$.
11. Prepare 0.1% cresyl-violet acetate (Sigma Aldrich) in deionized water by mixing 250 mg cresyl violet into 250 mL DI $H_2O$
12. After at least 5 min in DI $H_2O$, move slides into cresyl violet solution for 3 min
13. Dip slides 5 times in DI $H_2O$
14. Complete an ethanol dehydration sequence: place slides into 25% ethanol for 1 minute, 50% ethanol for 1 minute, 75% ethanol for 1 minute, 90% ethanol for 1 minute, 95% ethanol for 1 minute, 100% ethanol 2x for 2 minutes, Xylenes 2x for 2 minutes (can stay in xylenes indefinitely)
15. Take slides out of Xylenes 1 at a time, drop on 3 drops of Fisher Chemical Permount Mounting Medium (ThermoFisher Scientific, Waltham, MA, USA) and coverslip. Allow to dry flat overnight

### SM3. Calculations of space use and homing descriptors

Cumulative movement (meters)

Cumulative movement was calculated by adding the distances of all displacements between observations of a toad, as provided in the "ltraj" object created in the 'adehabitatLT' package (in meters). Cumulative movement was calculated separately for baseline movements and for movements following translocation.

Daily movement (meters/day)

Daily movement was calculated by dividing the cumulative movement observed for each individual during the baseline tracking period by the duration of this period, creating a daily movement metric.

Movement range (meters)

Movement range was calculated by finding the maximum distance between any two points for an individual during the baseline tracking period.

Straightness index

Straightness index of the homing path was calculated by dividing the length of the cumulative path of return by the straight-line distance from the release point to the return point. A straightness index of 1 indicates a completely straight path while index values approaching 0 indicate non-directional movement.

Duration of homing (hours)

Duration of homing was calculated by measuring the time from release to the point at which the toad first reached a point within 200 meters of one of its baseline points.

Homing speed (meters/hour)

Homing speed was calculated by dividing the distance the toad moved in the homeward direction (not the cumulative distance moved) by the homing duration.

Active homing duration/speed(hours and meters/hour)

To account for stationary periods following translocation, we also calculated "active homing" duration and speed. Active homing duration was calculated by subtracting the time it took the toad to move at least 100 m in the home direction from the total homing duration; active homing speed was calculated by dividing the homeward distance left to move following this initial movement of at least 100 m by the active homing duration.

Similarity of movement range before and after ablation treatment

To estimate the similarity of a toad's home range before and after ablation treatment, we plotted minimum convex polygons in ArcGIS Pro encapsulating all location points pre-ablation and calculated the proportion of post-ablation relocations that fell inside the polygon.

**SM4. Statistical treatment of toad movement data**

Normality was verified using the Shapiro-Wilks Test. Toad size (SVL) was compared between sexes using a Student's t-test and between field sites by ANOVA with a post-hoc Tukey’s test. Comparisons of mass, movement range, and homing duration/speed between field sites were done with Kruskal-Wallis tests with a post-hoc Dunn’s test. Wilcoxon Rank Sum Tests were used to compare mass and movement range between sexes. It was also used to compare the effects of translocation distance on homing descriptors (straightness, duration, speed, and time to move 100 meters homeward), as well as differences between sham and ablated individuals in home area pre/post manipulation, homing duration, and cumulative movement. A Paired Wilcoxon test was used to compare movement before and after ablations for sensory manipulated toads. Pearson’s Chi-Squared tests were used to compare binary homing success between field sites, translocation distances, sex, and sham/ablated individuals. The effect of toad size on movement range and homing duration was modeled with a linear model using the ‘lm’ function in the ‘stats’ R package [8] and with a linear mixed-effects model using the ‘lme4’ R package [9], where either movement range or duration was the response variable, size was the fixed effect, and field site was the random effect.

We used the ‘circular’ package in R [10] to analyze the directionality of movement angles between homing and non-homing toads. The “ltraj” objects generated using the ‘adehabitatLT’ package provide absolute angles (i.e. compass direction) and relative angles (i.e. the change in angle from the previous movement) for each movement segment. In our comparisons, we calculated the circular mean of all absolute angles of movement for each toad and normalized with respect to the home direction. We used the Rayleigh test of circular uniformity to test whether mean absolute angles differed from the home direction in homing and non-homing toads. We also compared the distribution of mean angles directly between homing and non-homing animals using a MANOVA based on trigonometric functions [11] using ‘manova’ functions in the ‘stats’ R package [8]. We also used the Rayleigh test to determine whether direction of translocation was evenly distributed.

**SM5. Statistical treatment of cell count data**

We used generalized linear mixed models (‘glmmTMB’ in R [12]) to test for differences in brain activity and its relation with behavior, with homing condition, brain region, and their interactions as the main predictors. Models used a negative binomial distribution and best model fit was confirmed using the ‘DHARma’ package in R [13]. Homing condition, brain region, and their interaction were the main effects predicting the number of pS6-positive cells. Initially, sex was included as a predictor. However, as sex yielded no significant effects on active cells and we saw no differences in movement range or homing performance due to sex, it was excluded as a predictor in the final models. Toad ID was included as a random factor and brain region area was included as an offset variable to account for brain area size. We modeled differences in brain activity due to ablations in a separate model where baseline animals were excluded, using the same model parameters as above except that ablation was also included as a main effect. Post hoc pairwise contrasts (between homing condition groups and ablation treatments) were calculated with estimated marginal means using the “emmeans” package in R [10].

To test if brain activity was related to either straightness or duration of homing, we performed linear mixed effects models using the “lme4” package, where pS6 count was the response variable and the homing descriptor (straightness or duration), brain region, and their interaction were fixed effects. Toad ID was a random effect and brain region area was an offset variable in the model. Linear mixed effects models were also performed within count data specific to each brain region, in which the response variable was pS6 counts within a given region, the homing descriptor was the fixed effect, Toad ID was a random effect, and region area was an offset variable.
